# Immuno-proteomic interrogation of dengue infection reveals novel HLA haplotype-specific MHC-I antigens

**DOI:** 10.1101/471821

**Authors:** Kavya Swaminathan, Niclas Olsson, Peder J. Lund, Caleb D. Marceau, Lisa E. Wagar, Yuan Tian, John Sidney, Daniela Weiskopf, Karim Majzoub, Aruna D. de Silva, Eva Harris, Mark M. Davis, Alessandro Sette, Jan E. Carette, Joshua E. Elias

## Abstract

Broadly effective vaccines against dengue virus (DENV) infection have remained elusive, despite rising infection rates in the developing world. Infection-specific peptide ligands presented on Major Histocompatibility Complexes (MHC) open new avenues for developing T-cell-based interventions. Past efforts towards mapping viral MHC epitopes were based on computational predictions that only partially reflected actual antigen presentation. To empirically identify DENV-specific MHC ligands, we developed an immuno-proteomics approach for interrogating DENV- and self-derived MHC ligands from infected B-lymphocytes. Here, we report four fundamental findings: First, over 700 infection-specific MHC-ligands reflected host cellular responses to DENV that were not apparent from the proteome. Second, we report 121 viral MHC-I ligands (108 novel) which clustered into discrete hotspots across the DENV polyprotein, some of which spanned DENV polyprotein components, described here as MHC ligands for the first time. Third, we found DENV ligands which were distinctly presented by MHC alleles previously associated with either high or low anti-DENV response. Fourth, we demonstrate that while our *in vitro* assay only overlapped with a small fraction of previously described DENV T-cell epitopes, several novel MHC ligands identified here were recognized by T-cells from DENV-infected patients despite having low binding affinities. Together, these discoveries suggest that virus and host-derived MHC ligands have under-exploited potential for describing the cell biology of DENV infection, and as candidates for designing effective DENV vaccines.

## Introduction

As the most widespread mosquito-borne viral disease, DENV infection is responsible for over 100 million estimated annual cases worldwide and is a leading cause of hospitalization and death in the developing world^1^. Rising DENV infection rates are particularly alarming since broadly effective vaccines and antiviral therapies have remained elusive. Traditional vaccine development approaches have focused on eliciting humoral immune responses against surface-exposed DENV antigens, notably the viral envelope (E) and membrane (M) proteins. However, humoral responses to vaccines and DENV serotypes 1-4 have been inconsistent, cross-reactive, and may promote antibody-dependent enhancement (ADE) of infection^2^. Accordingly, several vaccination trials were halted for exacerbating disease severity in patients, especially children^3-5^. Vaccines targeting T-cells could overcome limitations of strictly humoral antiviral responses^6,7^.

Unlike antibodies, which only bind surface-exposed proteins, cytotoxic CD8+ T-cells can theoretically target any viral protein if its immunogenic epitopes are presented by class I major histocompatibility complex (MHC-I) the surface of infected cells. The internal non-structural (NS) viral proteins NS1, NS3, NS4A, NS4B, and NS5 contain over 95% of the highly conserved regions across the four DENV serotypes (>80% sequence identity)^8^, while conversely, the remaining proteins (e.g. E and M) account for <5% of the highly conserved regions^8^. T-cell-mediated responses against the NS viral proteins^9^, therefore, stand to be broadly effective and circumvent ADE. Furthermore, since peptide ligands presented by MHC (pMHCs) report a cell’s internal status to the immune system^10^, their unbiased experimental interrogation could reveal new mechanisms by which cells respond to viral infection. This information can, in turn, be harnessed to identify host and viral biomarkers and vaccine targets.

Defining infection-specific pMHCs is an important preliminary step for developing prophylactic or therapeutic T-cell responses. This has often been carried out by first predicting likely high-affinity MHC-interacting peptides *in silico*^11^ and then testing them *in vitro* with high-throughput MHC binding^12^ and T-cell reactivity assays^13-15^. This strategy has revealed thousands of self- and pathogen-derived peptides capable of T-cell stimulation^16^. However, peptides eluted from MHC complexes and empirically identified through high-throughput mass spectrometry (MS) suggest that pMHC presentation may extend beyond MHC-ligands with high binding affinity^17^. Regardless of their affinities, *bona fide* pMHC may not elicit robust T-cell responses *in vivo*^18^. Thus the critical relationship between binding affinity, presentation propensities, and immunogenicity is incompletely captured by any single computational or experimental discovery approach alone. Empirical evidence combining multiple domains stands to bridge the gap between predicted candidates and those that are prophylactically relevant. This notion led to several recent studies which empirically examined antigen presentation from *in vitro* infection models of VACV, HIV, HRSV, HCV, and HPV^19-24^. These kinds of studies stand to reveal new T-cell targets and self-antigens which complement traditional epitope discovery approaches.

Here, we present an empirical immunoproteomic investigation of MHC-I antigens induced by DENV infection. By directly assaying MHC-I ligands presented by DENV infected cells we uncovered over 745 infection-specific viral and host-derived pMHCs. These include more than 100 previously unreported DENV peptides: most are predicted to bind MHC with low affinity. We note previously uncharacterized “junction-peptides” spanning multiple DENV proteins. We also describe distinct pMHC repertoires presented by different HLA alleles, complementing previously observed differences in T-cell responses. By comparing empirically isolated DENV pMHCs and those discovered by predictionbased methods, we demonstrate a complex relationship between binding affinity, presentation propensity, and T-cell recognition. Notably, the pMHC we identified were almost entirely distinct from previously described DENV epitopes^16^. We further show that CD8+ T-cell recognition or responses measured from DENV-exposed patients support five new DENV epitopes. Two of these five were highly conserved across all four DENV serotypes and two have poor MHC binding affinities. These findings suggest that conserved low-affinity epitopes could be promising candidates for broad acting DENV T-cell vaccines which span multiple DENV serotypes.

## Results

### DENV infection modestly perturbs the host proteome

To study how DENV infection modulates MHC-I antigen presentation we used a B-lymphocyte cells line (Raji; HLA-A*03:01/A*03:01, HLA-B*15:10/ B*15:10, HLA-C*03:04/C*04:01) engineered to exogenously express the receptor DC-SIGN^25^ widely used as models to study DENV infection *in vitro*^25-27^. We subsequently infected these cells with DENV serotype-2 (**Supplementary Figure 1**).

Infection-specific ligands could arise simply from induced protein expression. Alternatively, such ligands could result from virally-modulated antigen presentation pathways within infected cells, even if the underlying antigen proteins’ expression remains unchanged. We first compared protein levels before (control) and after infection to distinguish these two possibilities. We performed multiplexed proteome quantitation to compare control and DENV-infected Raji cells in duplicate (**Figure 1a**). Based on the 5,361 host and viral proteins quantified from control and DENV2-infected cells (**Supplementary Table 1**), we found that DENV infection only modestly perturbed the cellular proteome. As expected, DENV polyprotein components were the most distinguishing feature we identified from infected cells (fold-change>17; p<0.0001; **Figure 1b, Table 1**). However, <1% of human proteins (33) demonstrated significantly and substantially altered expression following infection (t-test, p<0.01, |fold-change|>2; **Figure 1b, Table 1**). Of these, four were previously reported to interact directly with the DENV polyprotein (**Table 1**). Signal transducer and activator of transcription 2 (STAT2), for example, decreased almost four-fold following infection, consistent with previous reports that immature DENV RNA polymerase NS5 protein induces its degradation through direct binding and subsequent ubiquitylation and proteolysis^28^. Six other proteins are known to be modulated by cellular pathways following DENV infection (**Table 1**). For example, sphingomyelin synthase 1 (SGMS1, fold change=5.1) was shown to facilitate viral attachment and infection of flaviviruses^29,30^ and its increased abundance in infected cells is consistent with DENV’s known role in interfering with lipid homeostasis^31,32^. We also noted the transcription regulator Zinc finger protein 729 (ZNF729) was the most upregulated host protein we measured (fold-change>19). Although this gene is poorly characterized, its location within a cluster of endogenous retrovirus-responsive zinc finger genes on chromosome 19^33^ could reflect its possible role in the innate antiviral response.

**Figure 1.**
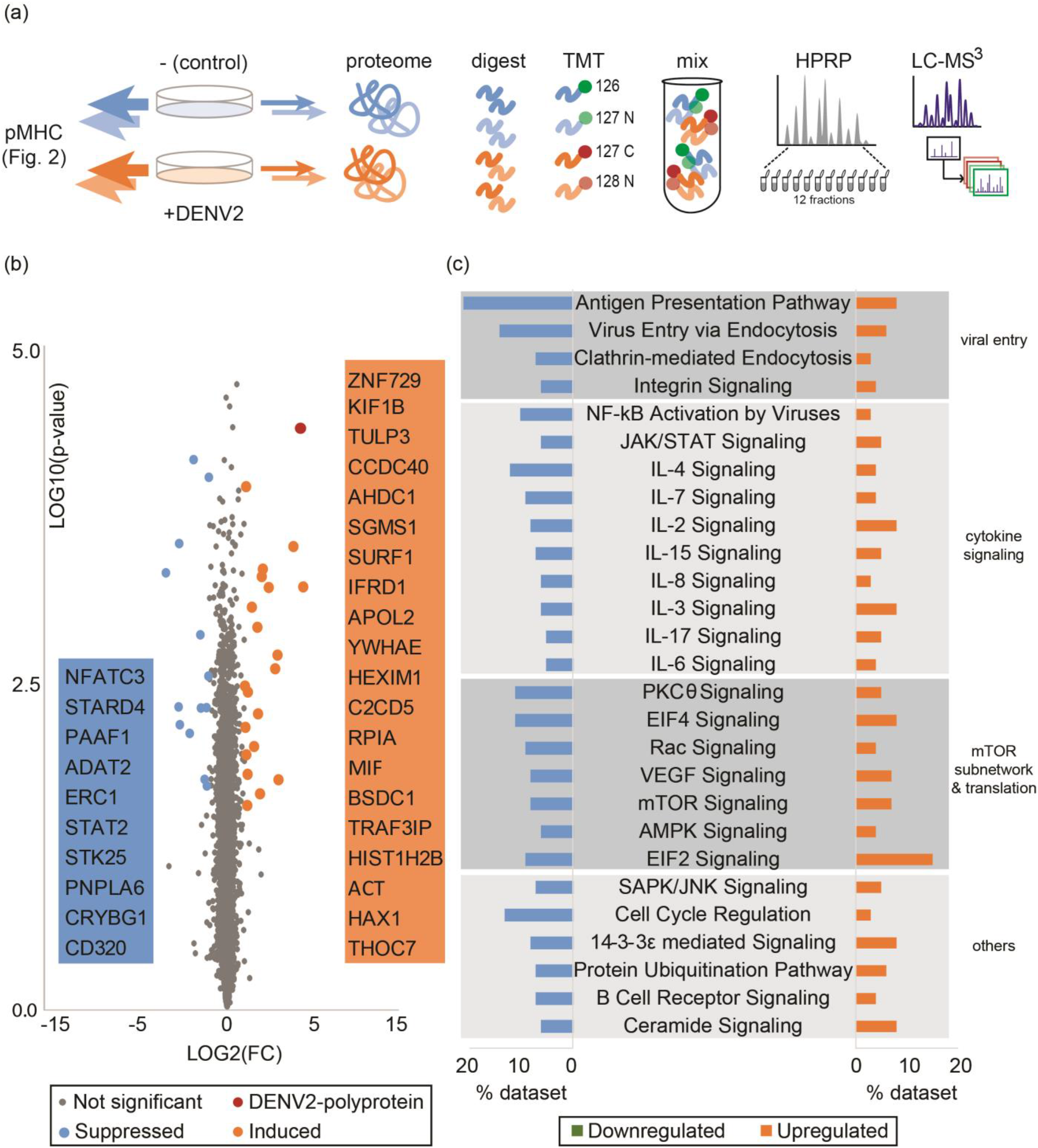
DENV infection modestly affects cellular proteome. **a)** Proteomic workflow. Duplicate cultures of DC-SIGN-expressing Raji cells were infected with DENV-serotype-2 (MOI = 5) harvested 27 hpi along with uninfected control cells. Most (98%) of the resulting lysate was used for the experiments described in Figure 2; the remainder was digested with trypsin, labeled with Tandem Mass Tags^59^, fractionated by concatenated high-pH reversed phase chromatography^60^, and analyzed by LC-MS with the multi-notch MS3 strategy^61,62^. **b)** Volcano plot comparing fold change in abundance (x-axis; Log2 FC) to the p-value (y-axis; t-test). Proteins significantly (p < 0.01) induced (Log2 FC > 1) or suppressed (Log2 FC < 1) during DENV infection are highlighted in orange and blue respectively. The DENV polyprotein (DENV2-polyprotein) is highlighted in red. Significantly changing host protein gene symbols are noted in decreasing order of absolute fold-change. **c)** Key pathways significantly (p < 0.001) modulated during DENV infection were inferred using Ingenuity Pathway Analysis of the 1003 proteins that were significantly (p < 0.01) changed upon infection. Pathways were categorized based on cell signaling function and are indicated on the right. The horizontal axis reports the percent of proteins assigned to the indicated pathways which were upregulated (orange) or downregulated (blue).

**Table 1.**
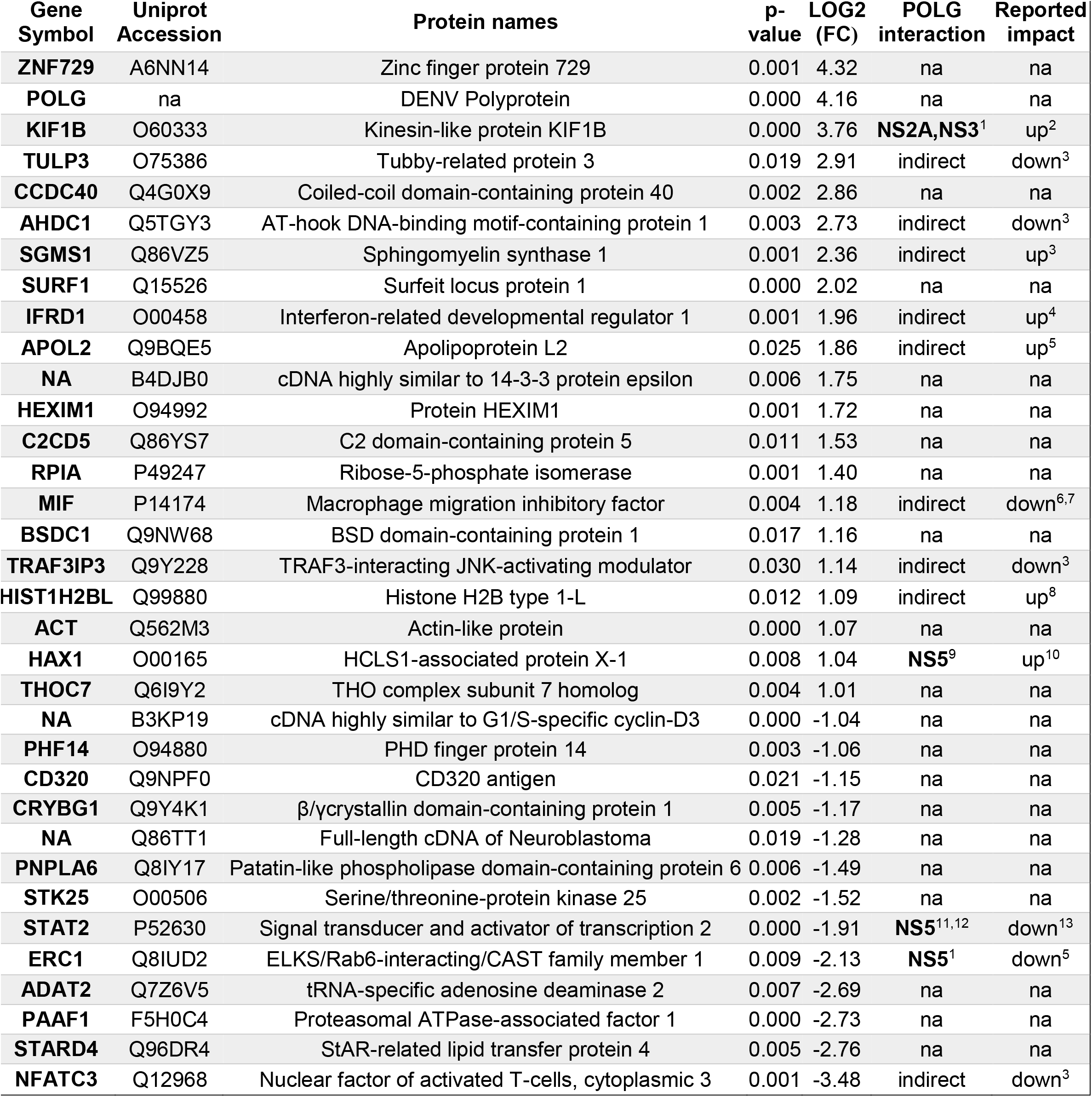
Proteins significantly and substantially modulated by DENV infection. Proteins with significant (p < 0.01, t-test) and substantially (|Log2 FC| > 1) increased or decreased abundance following DENV infection. For each entry, gene symbols, UniProt accessions, p-values (t-test), Log2 fold change of abundance upon infection, any known direct interactions with the DENV polyprotein (POLG) and any recorded impact on its abundance at a genome, transcriptome or proteome level upon DENV infection are noted.

The modest proteome-wide changes we measured upon infection led us to consider whether smaller-magnitude yet statistically significant changes in protein abundance reflected cell responses to DENV infection. We found that cellular processes such as translation, degradation, antigen presentation, and viral responsiveness were significantly enriched among 1,003 proteins that were significantly (p<0.03, q<0.2), though not substantially (|fold-change|<2), changed upon DENV infection (**Figure 1c**). Each of these categories could directly influence the quantity and quality of antigens presented by MHC. This led us to predict that DENV-altered pMHC repertoires could extend well beyond the 33 proteins showing the most changed abundances.

### Cellular response to DENV infection reshapes pMHC repertoires

Although thousands of DENV-encoded peptides have been tested for MHC binding and immunogenicity^16^, the extent to which DENV infection alters host pMHC repertoires remains unclear. Using the immuno-proteomic approach outlined in **Figure 2a** we mapped 5,397 unique pMHCs across all experimental conditions (**Figure 2b**). These include 745 pMHCs only eluted from DENV-infected cells, 71 of which were derived from the DENV polyprotein itself (**Supplementary Table 2a**). These peptides strongly conformed to the expected 9-mer-biased length distribution (**Supplementary Figure 2a**) expected for MHC-I ligands, supporting the validity of our dataset.

**Figure 2.**
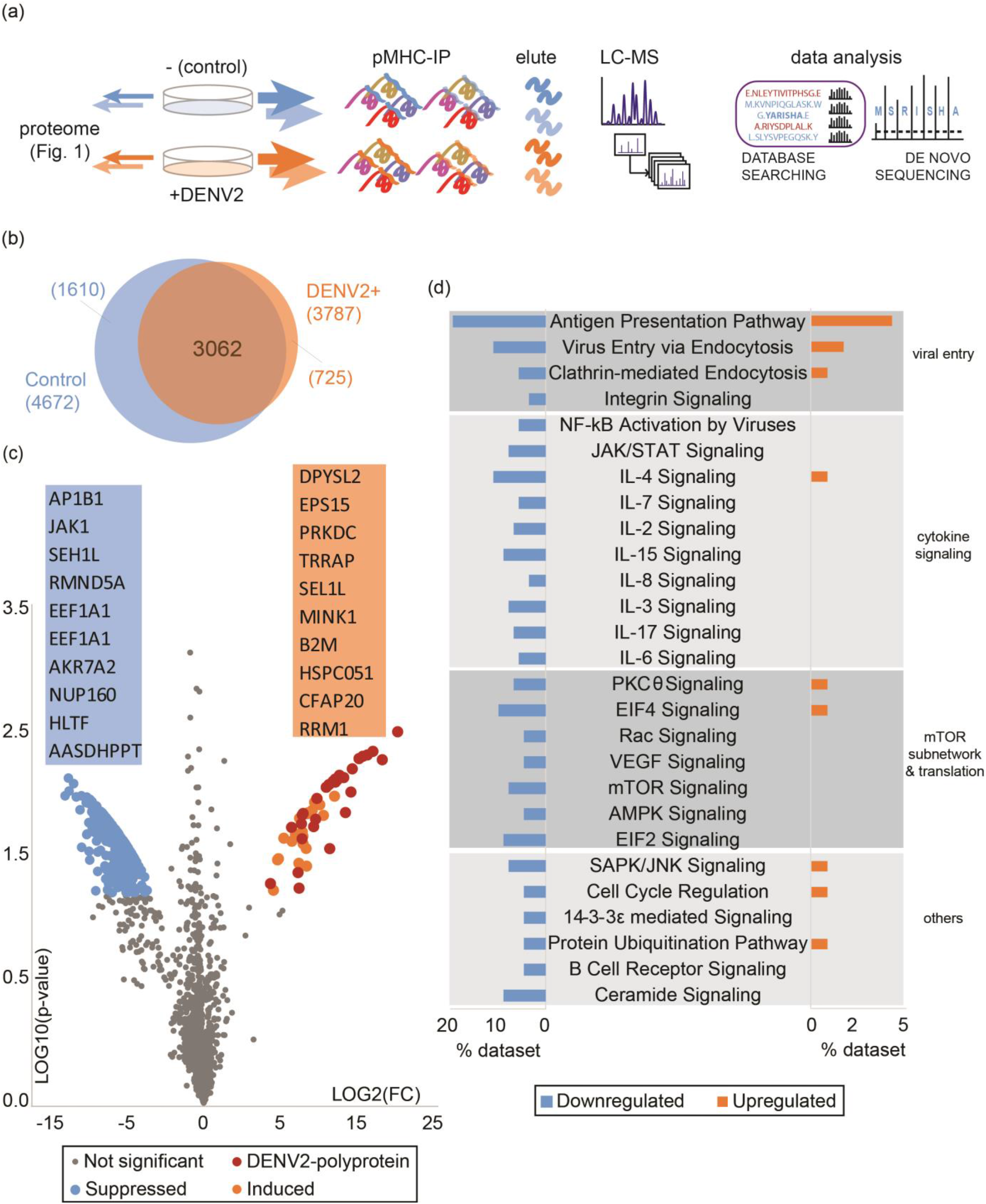
DENV infection has a strong influence on pMHC repertoires. **a)** Schematic summarizing the direct survey of antigen presentation during viral infection; peptide-MHC complexes were immunoprecipitated from control and DENV infected cells 27hpi. pMHCs were eluted off, desalted, and analyzed by LC-MS/MS. pMHC sequences were inferred using a combination of database searching (SEQUEST, PEAKS 7.5) and *de novo* sequencing (PEAKS 7.5) strategies. **b)** Venn diagram depicting the overlap between the pMHC repertoire of control (blue) and DENV infected (DENV2+) cells (orange); **c)** Volcano plot of fold change in pMHC relative abundance (Log2 FC) versus the p-value (t-test). 1,729 pMHCs consistently across both biological replicates of either control or DENV-infected cells (Supplementary Figure 3a) were used for this comparison. pMHCs significantly (p<0.01) suppressed (Log2 FC < 6) or induced (Log2 FC > 6) during DENV infection are highlighted in blue and orange respectively. DENV-derived pMHCs (DENV2-polyprotein) are marked in red. Gene symbols corresponding to the top ten pMHCs that increased and decreased upon infection are noted in decreasing order of absolute fold-change. **d)** Changes at the pMHC level in pathways from Figure 1c as inferred from 347 significantly (p<0.01) changed pMHCs upon infection. Percent of pMHC source proteins found in the dataset as being upregulated (orange) or downregulated (blue) from pathways are indicated on the horizontal axis. Pathways were categorized based on cell signaling function and are indicated on the right.

We quantified pMHC relative abundances using label-free quantification based on MS peak area of peptide precursor. Our data confirm this established approach ^34-37^ affords deep and reproducible survey of pMHCs (**Supplementary Figure 3a**) and robust comparisons between multiple immunopeptidomes (**Supplementary Figure 3b**). We found pMHC-I repertoires decreased in diversity following infection, with 18.9% fewer unique pMHC-I in DENV-infected Raji cells relative to uninfected controls (**Figure 2b**) (control: 4,672 total, x̅=3526 ± 456 per replicate; infected: 3,787 total, x̅=2497 ± 671 per replicate; p<0.001, Wilcoxon Signed Rank test, n=2). This observation could result from decreased MHC-expression, as has been reported for HIV, CMV and KSHV^38-40^. However, we found that MHC-I levels, as measured by proteome quantification (fold-change~0.96 - 0.98; p>0.01) and by flow cytometry, were not substantially changed between control and infected states (**Supplementary Figure 2b-c**).

Alternatively, a small number of highly abundant pMHCs (viral or self) could outcompete the basal self-repertoire for MHC-binding or detection. To test this, we ranked all pMHC by their relative abundances (see methods) with respect to each control or infected data set (**Supplementary Figure 3b**). Most pMHC we measured showed fairly consistent levels, with >87% contained within the first quartile of abundance rank deviations between the infected and control datasets (**Supplementary Figure 3c**). Since viral pMHCs were among the most abundant peptides we measured from infected cells, their presentation could account for a displaced sizeable proportion of self-peptides: more than one-third of pMHCs identified in the control state were not identified from DENV-infected cells (**Figure 2b**). Furthermore, most (>60%) of these were in the lowest quartile of abundance (**Supplementary Figure 3c**) despite their strong predicted binding affinities (**Supplementary Figure 3d**). This loss in sensitivity could be biological or technical, stemming from viral peptides outcompeting low abundance self-peptides for MHC presentation or for MS detection, respectively. It is difficult to distinguish these two scenarios from our data. However, we found no evidence of these “missing” peptides’ precursor ions from the infected datasets’ raw data files (data not shown). This supports a biological, rather than technical explanation for the differences between these data sets. Either explanation, however, supports the hypothesis that virus-induced pMHCs cause qualitative and quantitative shifts in self-pMHC presentation.

This finding further supports an incongruity between protein expression and antigen presentation. We measured ten-fold more pMHC which decreased in relative abundance following DENV infection than those that increased (**Figure 2c**), whereas two-fold more proteins increased in abundance following infection (20/33) than decreased (**Figure 1c**). We found no correlation between the magnitude of infection-induced pMHC changes and the corresponding changes in source protein abundance (R^2^<0.005) (**Supplementary Figure 4**). We found just one of the twenty upregulated proteins – Ribose-5-phosphate isomerase (RPIA) – was represented in the pMHC repertoire at all, but its pMHC relative abundance decreased 2-fold in the infected cells (**Supplementary Figure 4**). Conversely, of 13 proteins with decreased expression following DENV infection (**Table 1**), we identified three peptides with increased representation in the pMHC repertoire: These include peptides GHFEKPLFL - derived from patatin-like phospholipase domaincontaining protein (PNPLA6), RIYFRLRNK, and SLSPVILIK derived from beta/gamma crystallin domain-containing protein (CRYBG1) (**Supplementary Figure 4**). While these proteins have no reported roles in DENV infection, decreased PNPLA levels may be a result of DENV’s interference with cellular lipid metabolism^31,32^. Together, these data suggest protein expression changes do not directly predict novel antigen presentation in this system. Instead, they support a model in which pMHC repertoire alterations integrate multiple pathway-level changes within infected cells.

We further explored this notion by evaluating the functional pathway categories which changed following infection, comparing the pMHC and proteome datasets. We found several signaling pathways were implicated by both proteome and pMHC data sets following infection, including EIF2/4, mTOR, viral entry and antigen presentation pathways (p-value<0.03, q<0.2) (**Figure 1c**, **Figure 2c**). This suggests these pathways may be primarily responsible for the distinct pMHC repertoires we observed from DENV infected cells – both in the component proteins of these pathways, as well as the ultimate pathway targets. More generally, these data indicate that pMHC repertoires reveal infection-specific cellular responses which may not be as apparent from protein expression alone.

### DENV pMHCs tend to be restricted to discrete polyprotein “hot spots”

DENV-derived pMHCs are obvious clinical target candidates since they share little sequence homology with host proteins. Accordingly, over 800 predicted and empirically assayed DENV epitopes have been cataloged and represented in the Immune Epitope Database (IEDB)^16^. Comparing the DENV ligandome presented here with the IEDB resource allowed us to evaluate the relationships between peptide-MHC-binding, *in vitro* MHC-I presentation, and peptide immunogenicity.

Peptides derived from the DENV polyprotein were the most abundant pMHC class we identified in our dataset. Just 71 unique viral pMHCs (**Supplementary Table 3**) accounted for 4.6% of the total pMHC relative abundance (>3700 peptides) measured from DENV-infected cells, while viral polyprotein abundance in the proteome only accounted for 0.03% of the total cellular protein abundance (**Figure 2c**).

Interestingly, three DENV pMHCs – HRREKRSVALVPHVG, TAVTPSMTM, and ATMANEMGFLEK spanned cleavage sites between pr-M, M-E, and 2K-NS4B (respectively) within the polyprotein (**Figure 3a, Supplementary Table 3**). These pMHCs were reproducibly detected across replicate datasets and their lengths were consistent with other robust DENV pMHCs further substantiating their validity. pr-M spanning pMHCs could result from sampling immature virions, which are known to be abundant in infected cells^41,42^. However, M-E and 2K-NS4B junction sequences have not previously been characterized in model DENV systems. Our identification of pMHCs spanning this cleavage site suggests that the virus polyprotein could be sampled for presentation during active translation of viral mRNA and not only from mature polyprotein. Their predicted high binding affinity (<160 nM) to Raji endogenous alleles C*03 and A*03 respectively may further favor their presentation propensity.

**Figure 3.**
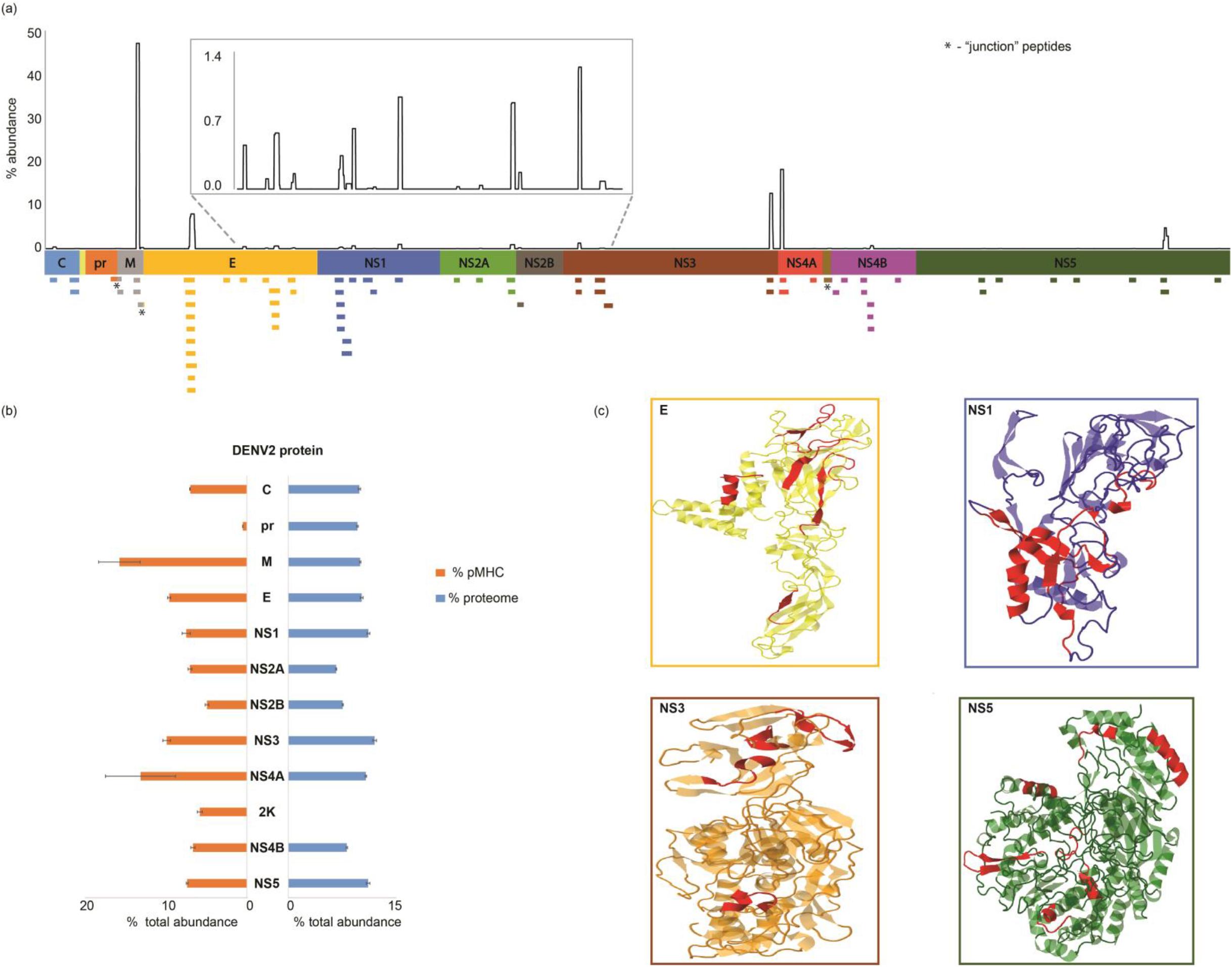
Biased pMHC presentation across the DENV polyprotein. **a)** Viral pMHCs isolated from DENV infected Raji cells mapped onto the viral polyprotein. Y-axis (top) represents relative abundance of pMHCs across the polyprotein. Individual unique pMHCs are indicated below. **b)** Summed peptide abundance of each polyprotein component in the proteome (blue) versus pMHC (orange). **c)** Secondary structure of pMHCs derived from E, NS1, NS3, and NS5 proteins are highlighted (red) as alpha helices or beta sheets on predicted tertiary structures using the Jmol platform. Wireframes represent unpredicted structures.

We found strong positional biases among the pMHC identified across the DENV polyprotein sequence. For example, a single peptide, THFQRALIF, derived from the DENV membrane protein M accounted for almost half of the total pMHC relative abundance measured across the DENV polyprotein (**Figure 3a**). We mapped other presentation “hotspots” across the E, NS1, and NS4A-4B proteins (**Figure 3a**). Such hotspots have previously been proposed to arise as a consequence of biases in cellular proteolysis or antigen processing machinery^43^. Alternatively, they have been suggested to be predictable based on measured HLA allele binding affinities^44^. We hypothesized that this presentation bias could also be influenced by intrinsic structural protein features.

### pMHC secondary structure influences DENV presentation hotspots

We first considered whether the positional biases we measured could be attributed to underlying differences in DENV protein abundances within infected cells. We found that DENV pMHC relative abundances across the polyprotein components did not mirror their source protein abundances (**R^2^ = 0.106, Supplementary Figure 5a**). Different protein or pMHC abundances measured across the polyprotein could also result from variable compatibility with our overall LC-MS workflow. For example, peptides derived from extremely hydrophobic proteins may have poor elution or ionization behavior, limiting their detection^45^ at both protein and pMHC levels (**Supplementary Figure 5b**). We found that protein hydropathies corresponded modestly (**Supplementary Figure 5c**) with measured proteome abundance (R^2^ = 0.554) while it had little association with pMHC hotspots we observed in **Figure 3a** (R^2^ = 0.002). We conclude that neither source protein abundance nor technical bias against hydrophobic pMHCs by the LC-MS explain the antigen presentation hotspots we observed across the DENV polyprotein.

It has been suggested that our immune systems evolved to present conserved viral protein features by MHC^46^. In support of this, we found that pMHC abundance showed some correspondence with the extent to which each DENV polypeptide component is conserved across homologous DENV protein sequences (R^2^ = 0.584) (**Supplementary Figure 5d**). This could further explain in part, the robust presentation of junction pMHCs that tend to be highly conserved (>78% identity). Selection pressures could act broadly on viral protein secondary structure to bias pMHC sampling. Accordingly, we found that alpha helices were significantly over-represented in the DENV pMHC repertoire compared to the virally derived peptides in the infected cellular proteome (**Figure 3c, Supplementary Figure 5e,f**). We attribute 60% of viral pMHC relative abundance and 45% of pMHC diversity to the presence of predicted alpha-helical secondary structures, while these regions accounted for just ~35% of the entire polyprotein (**Supplementary Figure 5e,f**). By comparison, predicted beta sheets, comprise ~21% of the polyprotein, account for 21.9% of the pMHC diversity and only 13% of the measured relative abundance (**Supplementary Figure 5e,f**). We also observed a general bias towards highly ordered alpha helix region being presented on MHC relative to the underlying proteome, and accordingly noticed a bias against the presentation of highly structured regions with low alpha helix propensities (**Supplementary Figure 5f**).

### HLA-B*35:01 restriction shifts positional bias towards presentation of NS protein pMHCs

Although our findings suggest protein secondary structure could influence DENV antigen presentation, HLA-haplotype-specific peptide affinities have a well-understood role in restricting the frequency and magnitude of *ex vivo* CD8+ T-cell responses^47,48^. Our dataset comprises pMHCs restricted by Raji cells’ endogenous alleles HLA-A*03:01, and HLA-B*15:10, which have been associated with weak CD8+ T-cell responses against DENV^47^. We hypothesized that HLAs associated with robust T-cell responses against DENV (e.g. HLA-B*35:01)^47^ could be effective because of the distinct pMHC repertoire they present.

To test this, we transduced Raji cells with FLAG-tagged HLA-B*35:01, and infected them with DENV. We then compared viral pMHC repertoires between parental cells lacking B*35, composite repertoires including B*35 from transduced cells, and pMHC repertoires attributed to B*35 alone (FLAG-immunoprecipitated). We found pMHC positional biases across the viral polyprotein which we associated with the B*35 allele were distinct from parental alleles (**Figure 4a**). For example, the M protein-derived peptide THFQRALIF accounted for half of the total viral pMHC relative abundance in parental cells, but this proportion dropped to just over 30% of viral pMHC relative abundance in the presence of B*35 (**Supplementary Figure 6a**). One explanation for this decrease is that the expanded pool of B*35 restricted pMHC competed with the cell’s endogenous alleles for MHC presentation. Accordingly, we found that levels of this peptide decreased to <2% of the total viral pMHC pool when we specifically measured B*35-bound pMHC (**Supplementary Figure 6a**).

**Figure 4.**
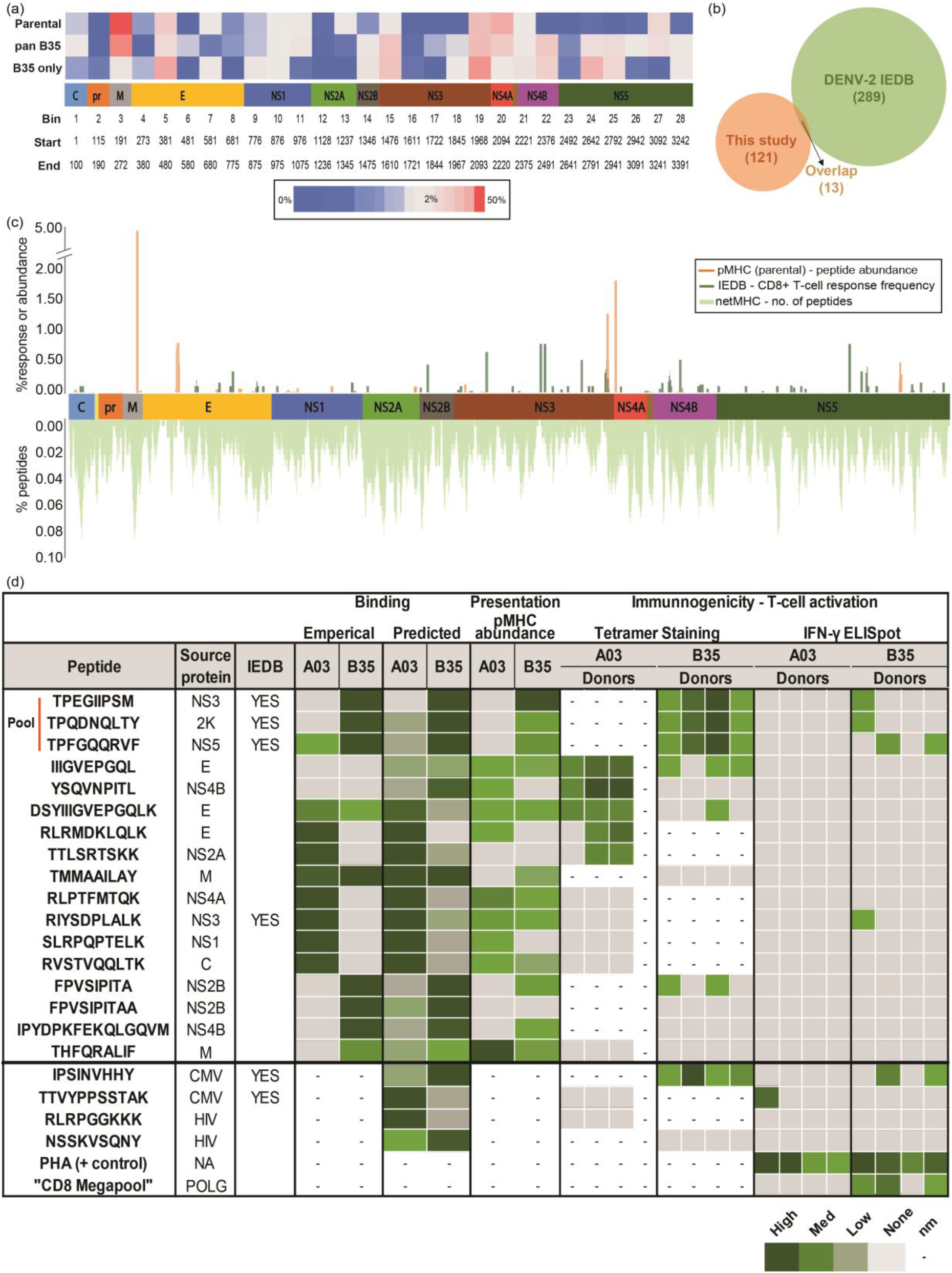
Host HLA shapes viral pMHC repertoire and T-cell response. **a)** Heatmap contrasting the presentation hotspots in Raji cells expressing endogenous HLA (bottom), pan-MHC repertoire of cells expressing HLA-B*35:01 (middle) and B*35-restricted pMHCs (top) across the polyprotein. Bins represent summed pMHC relative abundances across listed start and stop positions across the polyproteins. Protein sections with no pMHCs were not binned and polyprotein lengths were adjusted to reflect missing sections. Color scales represent the percentile ranks of summed pMHC relative abundances from each region within each repertoire. **b)** Venn diagram summarizing the overlap between all DENV pMHCs isolated in this study (orange) and the DENV2 epitopes (human host, positive assays) listed in the Immune Epitope Database (green). **c)** Distribution of endogenous Raji HLA restricted pMHC abundance (percentage of total viral pMHC) from this study (orange) versus IEDB epitope T-cell response frequencies (darkgreen) represented as percentages across the DENV polyprotein (x-axis). All 9-11 mer peptides predicted by netMHC to bind (< 5000 nM) to endogenous Raji HLAs (parental) are plotted below (light green) to reveal predicted binding hotspots. Y-axes represent the percentage of peptides deemed binding spanning any given residue. d) Contrasting binding affinity (predicted using netMHC and measured *in vitro*), presentation propensity (pMHC abundance from parental (A*03) and B*35-only (B35) experiments) and immunogenicity (tetramer staining and ELISpot) of seventeen DENV derived pMHCs across A*03 and B*35 HLAs. Tetramer staining and ELISpot assays were used to assess the frequency of T-cell responses against four A*03 and four B*35 positive donors. Phytohemagglutinin (PHA) was used as positive control in ELISpot assays. One A*03 and one B*35 restricted epitope each from CMV and HIV were used as positive and negative controls respectively in both assays. Frequency of response for three pooled IEDB peptides was divided equally. Color scales of frequencies in tetramer staining range from 0.01 (low) – 0.4% (high). Peptides were selected to balance their prior description in IEDB, their predicted binding affinities, source protein, and abundances as measured by LC-MS. nm = Not measured.

We measured fifty pMHCs (**Supplementary Figure 6a, Supplementary Table 3**) derived from thirty-seven regions along the DENV polyprotein that were not presented by parental cells. This pMHC repertoire shift manifested as increased E-protein presentation by B35-expressing cells: pMHCs derived from this protein increased 2.2 fold, from 9.6% to 20.99% of the total DENV pMHC repertoire we could specifically attribute to the B*35 (**Supplementary Figure 6b**). We also found substantially increased relative abundance and diversity of pMHCs derived from DENV non-structural (NS) proteins in B*35-expressing cells. This was most evident in the NS3 and NS5 proteins which demonstrated increased presentation levels (**Figure 4a, Supplementary Figure 6a**) as well as representation from distinct regions relative to the endogenous HLA alleles.

### Empirical pMHC presentation is partially predicted by MHC-binding affinity

The HLA allele-specific antigen presentation biases for NS and M proteins described above could most simply be explained by differences in binding affinities among putative peptide ligands. To address this, we predicted binding affinities for the endogenous Raji HLA alleles and B*35 across the DENV polyprotein sequence with the netMHC binding prediction algorithm^49^ (**Figure 4c, Supplementary Figure 6d**). We determined that the presentation biases described in **Figure 3** could not be clearly attributed to sequence motifs within the polyprotein: 9 – 11-mer peptides predicted to bind parental HLA alleles were widely distributed across all polyprotein components (**Supplementary Figure 6b-c**), covering 97% of all amino acid residues (**Figure 4a**). We also found that pMHC repertoires which changed in the presence of B*35 allele were not fully explained by predicted “B*35-binding hotspots” alone (**Supplementary Figure 6d**).

Each DENV protein sequence was predicted to yield peptides with at least modest affinities (<5000 nM) for the endogenous Raji HLA-alleles and for the exogenous B*35 allele (**Supplementary Figure 6d**). We therefore evaluated the extent to which binding affinity predictions corresponded with empirical pMHC measurements. This would allow us to evaluate the relationship between MHC-binding affinity – a key metric used for screening T-cell targets *in silico* – and eluted pMHC as measured by mass spectrometry. Considering the predicted MHC-binding affinities of 121 DENV pMHCs measured from parental and B*35-expressing cells, we found just over half were predicted to bind any of the Raji endogenous MHCs with even modest affinities (<5000 nM) (**Supplementary Table 3**). While the highly abundant pMHC (THFQRALIF) derived from the M protein corresponded with a predicted binding hotspot (**Figure 4c**), this particular peptide’s predicted binding affinity to endogenous Raji alleles was relatively low (a minimum of 1726 nM to B*15) (**Supplementary Table 3**). More generally, we found that DENV peptides predicted to bind HLAs with high affinity did not correlate with increased presentation abundance as measured by LC-MS (**Supplementary Figure 7a**).

Together, our observations suggested that predicted MHC-binding affinity only partially reflected empirically measured pMHC presentation. This led us to compare our DENV pMHC dataset with those previously reported in IEDB^16^. We focused this comparison on the subset of high-affinity DENV serotype-2 epitopes which were capable of T-cell stimulation *in vitro*. We found a stark incongruity between the DENV pMHCs we measured with our immuno-proteomic approach and those previously reported in IEDB. Only thirteen DENV serotype-2 pMHCs were shared between the 121 reported here and the 289 cataloged in IEDB (**Figure 4b**). Of these, eight were predicted to strongly bind (< 50nM) any of the Raji cells’ HLA alleles or B*35 (**Supplementary Table 3a**). We also found that the IEDB-catalogued epitopes, when restricted by Raji’s endogenous alleles, were more uniformly distributed across the polyprotein in proportion to protein length (R^2^ = 0.83) than our empirically determined pMHCs (R^2^ = 0.37) which were skewed towards a relatively small number of discrete sites (**Figure 4c, Supplementary Figure 7b-c**).

### Low-affinity DENV epitopes are recognized by patient-derived CD8+ T-cells

We observed a strong contrast between predicted MHC-binding and empirical presentation, measured from an *in vitro* culture system (**Figure 4b**). This highlighted the need to consider the influence both pMHC properties could have on T-cell immunogenicity. Although high-affinity MHC ligands are often prioritized for their ability to stimulate T-cell responses^50^, low-affinity pMHCs may be immunologically important due to other factors such as abundance, binding stability, and T-cell receptor (TCR) interactions^17,51,52^. Furthermore, antigens presented through our *in vitro* infection model may not precisely reflect those that are most commonly presented *in vivo*. To distinguish these possibilities, we synthesized seventeen DENV pMHCs and assessed their abilities to bind to two HLA alleles (A*03 and B*35) and to be recognized by and activate T-cells *ex vivo* (**Figure 4d**). Peptide-MHC affinities were predicted using the netMHC algorithm against parental Raji cell HLA alleles and HLA-B*35 and verified *in vitro* using competition assays. T-cell responses were measured with ELISpot or tetramer staining assays using PBMCs collected from DENV seropositive donors expressing either, but not both of the two alleles above. We found that empirical peptide-MHC binding measurements strongly agreed with predictions for all tested peptides, demonstrating the netMHC algorithm’s robustness. These binding predictions and measurements, however inconsistently matched the empirical immunopeptidome data we measured by LC-MS. For example, the peptide TPEGIIPSM from NS3 bound the B*35 allele with high affinity and was identified with high relative abundance. In contrast, the peptide TTLSRTSKK from NS2A bound the A*03 allele with similarly high affinity, yet was detected by LC-MS with very low relative abundance (**Figure 4d, Supplementary Table 3**).

Using MHC tetramer and ELISpot assays, we found that up to nine of seventeen peptides we tested were recognized by seropositive donors’ T-cells and/or trigger their IFN-γ production (**Figure 4d**). Three pMHCs previously described in IEDB as high-affinity B*35 restricted epitopes were pooled for tetramer staining assays, which indicated high frequency (0.06 - 1.2%) epitope-specific CD8+ T-cells (**Figure 4d, Supplementary Table 4**), and confirming their prior measurement. These data were further supported by the high response frequencies (> 0.1%) measured by companion ELISpot assays of each peptide in isolation. Surprisingly, we found at least two pMHCs with low predicted or measured binding affinity to A*03 (>5000 nM), yet small populations of T-cells (0.01 – 0.03%) were specific to these epitopes in all three A*03-donor samples we tested (**Figure 4d**). One of these pMHCs (IIIGVEPGQL) was derived from the DENV envelope (E) region [655 – 668], which was not previously characterized as a CD8+ T-cell target. One other pMHC (DSYIIIGVEPGQLK) with modest affinity (~2000 nM) was also derived from the same region of the E protein. The other pMHC, YSQVNPITL, derived from NS4B was also predicted to bind A*03 with low affinity but yielded strong tetramer staining. This peptide was predicted to bind B*35 with high affinity, but was neither identified in association with B*35 by LC-MS nor did it yield positive tetramer or ELISpot assays from any patient-derived specimens. We further note that this sequence is highly conserved (>90%, **Supplementary Table 3**) across all four DENV serotypes. Our LC-MS-based assay implicated another highly conserved (>80%) E protein epitope, RLRMDKLQLK, which was predicted to have high HLA-A*03 binding affinity (**Figure 4d, Supplementary Table 3**). However, we did not measure T-cells that recognized it, or eight other pMHCs predicted or measured to have high-(<50 nM) to moderate (<500 nM) binding affinity.

Upon testing PBMCs from both A*03 and B*35 donors, we found higher frequency responses to non-structural (NS) proteins than the structural proteins. Thus the shift we observed towards greater NS protein pMHC presentation by B*35 corresponded with the higher magnitude and frequency of anti-DENV T-cell responses previously observed in B*35+ individuals^47^. Furthermore, the M-protein derived pMHC, THFQRALIF found in high abundance in parental Raji cells but substantially decreased in B*35-Raji cells was a poor CD8+ T-cell target in our assays (**Figure 4d**).

The absence of correlation between both binding affinity, *in vitro* presentation and T-cell response (Supplementary Figure 7a, Table 4) support the notion that high-affinity binding or robust empirical presentation alone are insufficient to predict T-cell activation. However, our findings suggest that empirical pMHC presentation measured during infections provide an alternate set of epitopes than those discovered using bindingprediction based strategies and reveal previously unknown low-affinity antigens for the design of T-cell vaccines.

## Discussion

In this study, we considered multiple factors which could shape antigen presentation during DENV infection with the objective of identifying self and virally derived pMHC. Both the precise antigens and the processes that produce them could serve as foundations for developing new T-cell based interventions. Towards this end, we measured changes to the cellular proteome following infection as a way to understand how these changes might be reflected in infected cells’ MHC repertoires. We found that infection-induced modest protein-level changes but suggested key host-encoded antigen presentation pathways that were modulated directly or indirectly by DENV infections. Interestingly, we found that DENV protein abundances were not strictly equal given their simultaneous translation from a single mRNA. We predict that this could result in part from differential individual protein hydropathies that affect their MS measurement (**Supplementary Figure 5b-c**) but cannot rule out the impact of differential compartmentalization or half-lives in the cell.

While individual pMHCs showed little correlation with the host proteins from which they were derived (**Supplementary Figure 4**) we found that pMHCs reflected broad cellular pathway level changes. The discrepancy between source protein and pMHC abundances could result from virus-mediated changes to viral and host-protein synthesis or degradation, both of which could result in higher pMHC presentation^53,54^. Further studies combining protein turnover quantification and RNA-sequencing could clarify how the virus might alter antigen presentation through protein proteostasis mechanisms.

The contrast we observed between control and DENV infected pMHC repertoires also highlighted how antigen presentation can report internal cellular states for immune surveillance^54^. We found that pMHC repertoire changes provided insights into the cell biology of viral infection and immune response dynamics – insights that were not apparent from cellular proteome measurements alone. Our data further suggest that infectionspecific self-pMHCs could serve as attractive biomarkers of acutely affected cellular pathways during DENV infection. However, directing vaccine or antiviral therapies against self-pMHCs could be confounded by the expected lack of immune response to selfantigens. Additional studies comparing these data to other infection and cellular stress models could help resolve which self-pMHCs are truly DENV-specific.

Virus-derived pMHCs are intuitive targets for T-cell based interventions against DENV. Using our immuno-proteomic system, we found 121 DENV-derived pMHCs which were restricted by Raji cells’ endogenous alleles or the HLA-B*35:01 allele we transduced. This malleable approach allowed us to survey and compare the pMHC repertoire across HLA-alleles important for DENV infections. Across both reported low (e.g. A*03, B*15) and high (e.g. B*35) response alleles, we mapped presentation hotspots that were shaped at least in part by structural features of viral proteins. This supports the possibility that DENV proteins evolved to preferentially present some restricted domains within their proteomes to evade immune activation^46^. This is further supported by the positive correlation we observed between pMHC abundance and sequence conservation across DENV serotypes.

The most unexpected of viral pMHCs we observed from both parental and B35-Raji cells were four “junction peptides” derived from regions spanning DENV proteins. These pMHCs may result from the sampling of the actively translated DENV polyprotein. Alternatively, they could arise from mistranslated or erroneously spliced proteins broadly known as defective ribosomal products (DRiPs)^55^ as documented previously in other infection systems such as influenza^56^ and lymphocytic choriomeningitis viruses (LCMV)^57^. Either way, these highly conserved (78 – 97% identity across DENV serotypes) pMHCs two of which have high binding affinity to the Raji HLA or B35 alleles could be excellent candidates as T-cell targets.

We examined the role HLA-restriction has in shaping the viral pMHC repertoire by comparing the pMHC repertoire HLA alleles associated with low-grade DENV response (A*03, B*15) to pMHC restricted by the B*35 allele which was previously associated with high-grade DENV responses. We found sharply decreased M-protein presentation by B*35, but markedly increased presentation of several viral NS proteins. Our results indicated that these stark HLA-associated differences were not simply the consequence of additional B*35-preferred anchor residues in the NS proteins relative to the M protein. Notably, the M-protein derived pMHC, THFQRALIF found in high abundance in parental Raji cells but substantially decreased in B*35-Raji cells was a poor CD8+ T-cell target in our assays. This suggests that differences in anti-DENV response between A*03 and B*35 expressing individuals could be explained by non-immunogenic, yet highly abundant peptides (**Figure 4a**). Such peptides could outcompete more robust T-cell targets. Its low presentation during B*35-restriction could, therefore, facilitate better anti-DENV T-cell responses by presenting robust T-cell targets. We suggest that this observed shift towards increased NS protein presentation could underlie the stronger anti-DENV T-cell responses in HLA-B*35-expressing individuals^47^. However, difficulty obtaining PBMCs from individuals who expressed A*03 or B*35 but not both, and also had prior DENV serotype-2 exposure – limited our ability to test this finding with statistical power or to extend it to more DENV pMHCs or subjects.

We were initially surprised to observe that just 11% of the DENV pMHC ligands we identified here had prior positive evidence in *ex-vivo* screening assays from patient-derived T-cells^16^, and just 4% of previously described T-cell epitopes were confirmed as pMHC ligands here **(Figure 4b)**. We reasoned that this could result in part from only 58 (~20%) of IEDB epitopes being reported as restricted by Raji cells’ endogenous (A*03, B*15, C*03, C*04) or B*35 family of alleles. These two datasets’ discordance could also have technical causes. For example, *bona fide* T-cell epitopes may escape identification by LC-MS due to low abundance, incompatibility with the chromatography system, or poor ionization characteristics associated with epitopes we did not identify here. However, we identified self-peptides derived from an extremely wide dynamic range of protein abundances ranging from low-abundance transcription factors to highly abundant histones. Furthermore, DENV-derived peptides were among the most dominant features in our infected cell datasets (**Supplementary Tables 3, 4**), suggesting that low abundance is unlikely to be a sufficient explanation for the differences between these two datasets. Similarly, we did not observe significant biases for or against amino acid utilization within or between DENV- or self-MHC ligands, suggesting our assay’s ability to sample peptides with diverse physiochemical properties.

We instead attribute the incongruity we found between these datasets to fundamental differences in what their underlying assays measure: the elution methodology presented here measures peptides empirically found to be presented by MHC, without respect to T-cells or other cells of the immune system, and without a strict requirement for high binding affinity. By comparison, prior epitope screening procedures tested T-cells from multiple patients, each with TCR repertoires that could have been shaped by multiple prior DENV infections, and which most robustly responded to very high-affinity epitopes^50^. Thus, the peptides most frequently presented by the cells we infected *in vitro* may not precisely coincide with peptides that elicited strong immune responses across a wide range of patient conditions. Our finding that many of the DENV MHC ligands identified here were predicted to have low binding affinity supports this notion. Nevertheless, it is possible that such low-affinity peptides, when processed with high efficiency, could compensate for low binding affinity^17^. Some of these might also be immunogenic, as we demonstrate in **Figure 4d**. MS-based presentation evidence gives alternate peptide sets worth testing with expensive T-cell assays which may depend on very rare clinical specimens. Ultimately, however, further studies will be necessary to map out the complex relationships between antigen abundance, binding affinity, presentation propensity, and immunogenicity.

We believe our observation that several low binding affinity peptides were reproducibly recognized by CD8+ T-cells and/or stimulate IFN-γ production in seropositive individuals is particularly noteworthy. First, these results support our *in vitro* infection model as a reasonable proxy for reporting pMHC that are likely presented *in vivo*. Furthermore, since these peptides would not be identified through MHC-binding screens, they argue for improved prediction algorithms that incorporate pMHC attributes in addition to MHC-binding affinity^58^. Other factors such as protein abundance and promiscuous HLA-binding may play an important role in shaping antigen presentation^17,44^. As IEDB evolves to include MS-derived pMHCs, it will continue to be a valuable resource for developing these improved models.

The system we describe herein represents a method for the unbiased isolation of haplotype-specific MHC-I peptide antigens during DENV infection. While we cannot rule out any effects of HLA overexpression on antigen presentation, we note that similar systems have been used to successfully study HLA-haplotype specific antigen presentation^17^. The approach can be adapted to many other pathogens for a comprehensive survey of antigen presentation across multiple HLA alleles including those associated with resistance or susceptibility to disease. With it, researchers can isolate bonafide viral antigens that can be further vetted for immunological significance.

## Acknowledgments

The authors thank members of the Elias and Carette lab for discussions and feedback on the manuscript. We would also like to thank Dr. Angel Balmaseda and Cristhiam Cerpas at the Laboratorio Nacional de Virología, Centro Nacional de Diagnóstico y Referencia, Ministerio de Salud and Sustainable Sciences Institute in Managua, Nicaragua, for testing of samples and preparation of PBMCs from Nicaraguan blood bank donors. This work was funded by the W.M. Keck Foundation Medical Research Program (J.E.E.), the Damon Runyon Cancer Research Foundation (J.E.E.), and the Stanford Human Systems Immunology Center, supported by the Bill and Melinda Gates Foundation (J.E.E., J.E.C., and M.M.D.). NIH funding supported the work by grant P01AI10669 (E.H.) and contract HHSN27220140045C (A.S.).

## Author contributions

K.S. and J.E. conceived and designed the study and figures, and wrote the manuscript. K.S. carried out the experiments, compiled and analyzed the data and generated figures. N.O. performed pilot experiments. C.M. and K.M. cultured virus and generated HLA-transduced cell lines with guidance from J.C. P.L. designed HLA-constructs, carried out FACS and helped with data analysis. L. W. helped with multiplexed tetramer staining assay design, execution, and data analysis. M.D. provided guidance and feedback with the design of immunological experiments. Y.T., J.S., and D.W. performed peptide-MHC binding and ELISpot assays and analyzed the data with guidance from A.S. A.D and E.H provided Dengue seropositive PBMC samples.

## Declaration of Interests

The authors declare no competing interests.Figures

## Supplementary Resources

**Supplementary Figure 1.**
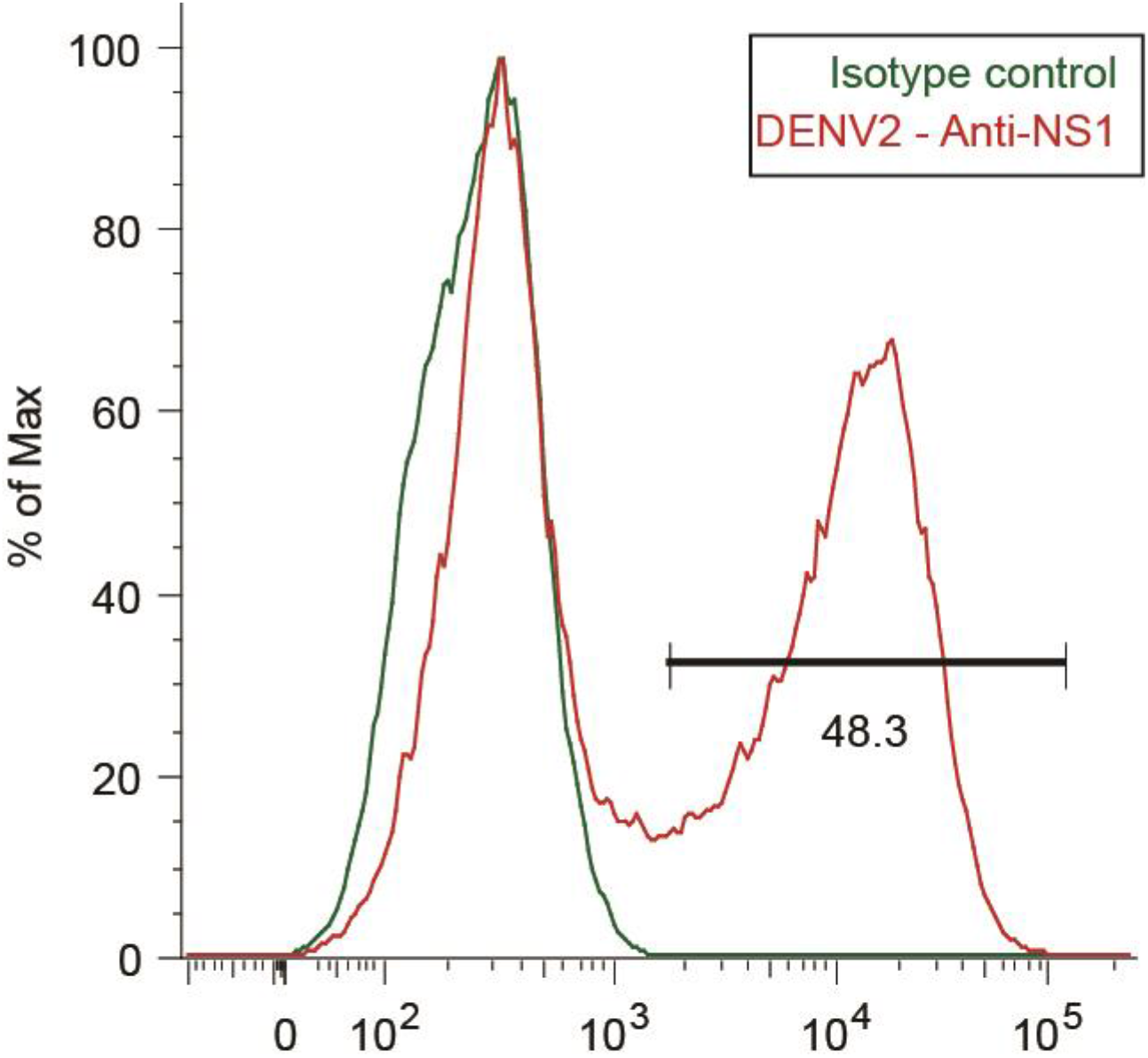
DC-SIGN-expressing Raji cells are competent for DENV infection. FACS traces of DENV2 NS1 (red) versus isotype control (green)

**Supplementary Figure 2.**
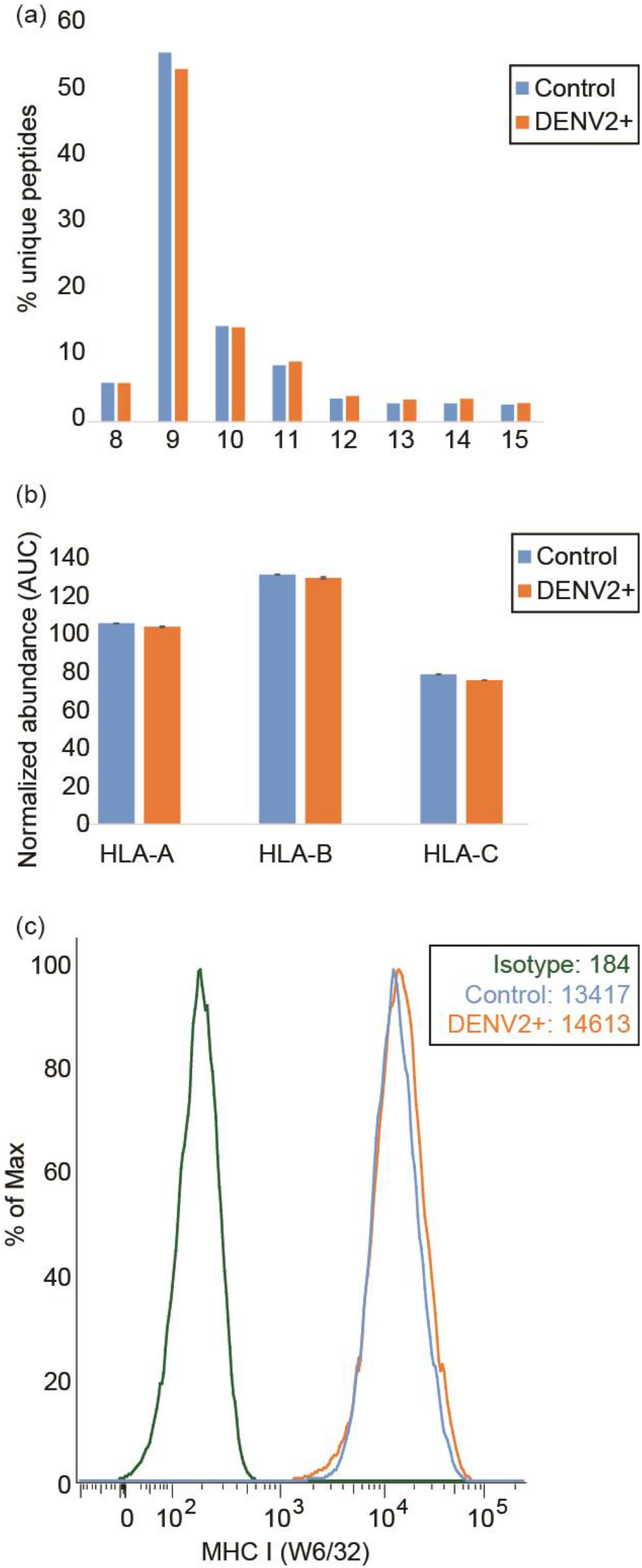
DENV infection has little impact on MHC-I peptide length and total MHC-I levels. **a)** Length distribution of pMHC repertoire in control (blue) and DENV infected – DENV2+ (orange) Raji cells. **b)** Summed abundance (normalized to total protein abundance) of tryptic peptides from MHC-I HLA-A, HLA-B, and HLA-C proteins in the proteome of control (blue) and DENV infected – DENV2+ (orange) cells. Error bars represent standard deviations across two biological replicates. c) FACS staining MHC-1 with the pan-MHC-I antibody (clone W6/32) in uninfected control (blue) and DENV infected Raji cells DENV2+ (orange). FACS trace for the isotype control is marked in green.

**Supplementary Figure 3.**
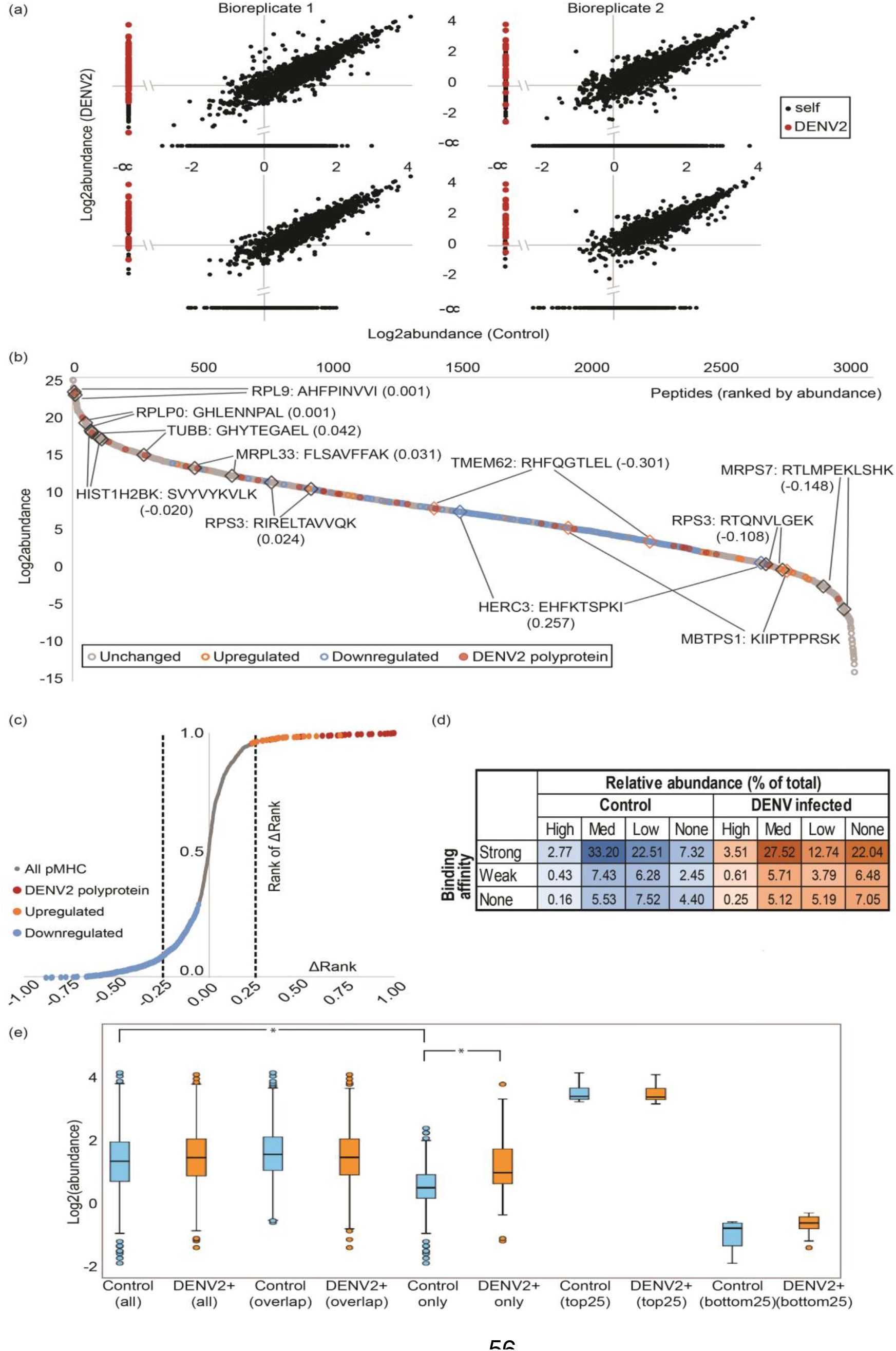
DENV derived pMHCs shift the endogenous pMHC pool. **a)** Log2 transformed pMHC relative abundance in control (x-axis) vs. DENV infected (DENV2+) cells (y-axis) across both bio-replicates. Unfiltered dataset plots are on top, bottom plots represent only pMHCs found in both bio-replicates. DENV pMHCs (DENV2) are highlighted in red. **b)** Distribution of log2 transformed abundance (y-axis) across pMHCs ranked in decreasing order of abundance (x-axis) in control and DENV infected cells. pMHCs whose rank changes were within the first quartile in grey, and those significantly up or down-regulated are shown in orange and blue respectively. Examples of each category - unchanged (black diamonds), up (orange diamonds) and down-regulated (blue diamonds) pMHCs in control and DENV infected datasets are highlighted and labeled with gene symbol of corresponding source proteins and the percentile change in abundance ranks following infection. **c)** Distribution of change in percentile rank (x-axis, ΔRank) in control and DENV infected cells colored according to the direction of fold change after infection. Y-axis represents the change in rank before and after infection calculated as a percentile rank - (Rank of ΔRank). DENV pMHCs (DENV2) are highlighted in red. **d)** Percentage of unique pMHC in each dataset categorized based on their predicted binding affinities to endogenous Raji alleles and their measured relative abundance in control (blue) and DENV infected (orange) datasets. pMHC were categorized based on their minimum binding rank (%) across the Raji HLA alleles calculated by netMHC as ‘strong’ (</= 0.5%), ‘weak’ (</=2%) or ‘none’ (>2%). Abundance levels of the pMHCs were deemed ‘high’ if they were in the top 25^th^ percentile, ‘med’ if they were between 25^th^ and 60^th^ percentile and ‘low’ if below the 60^th^ percentile. pMHCs not detected in a dataset were deemed ‘none’. **e)** Boxplots contrasting relative pMHC abundance (Log10 abundance) derived from control and DENV infected (DENV2+) Raji cells. Pairs of plots represent relative abundances in control and DENV infected datasets for all pMHCs; those identified in both control and infected states; pMHCs exclusive to either state; and the twenty-five highest and least abundance pMHCs. Pairs of datasets with significantly (p<0.001, Wilcoxon Signed Rank test, n=2) different means, are indicated (*).

**Supplementary Figure 4.**
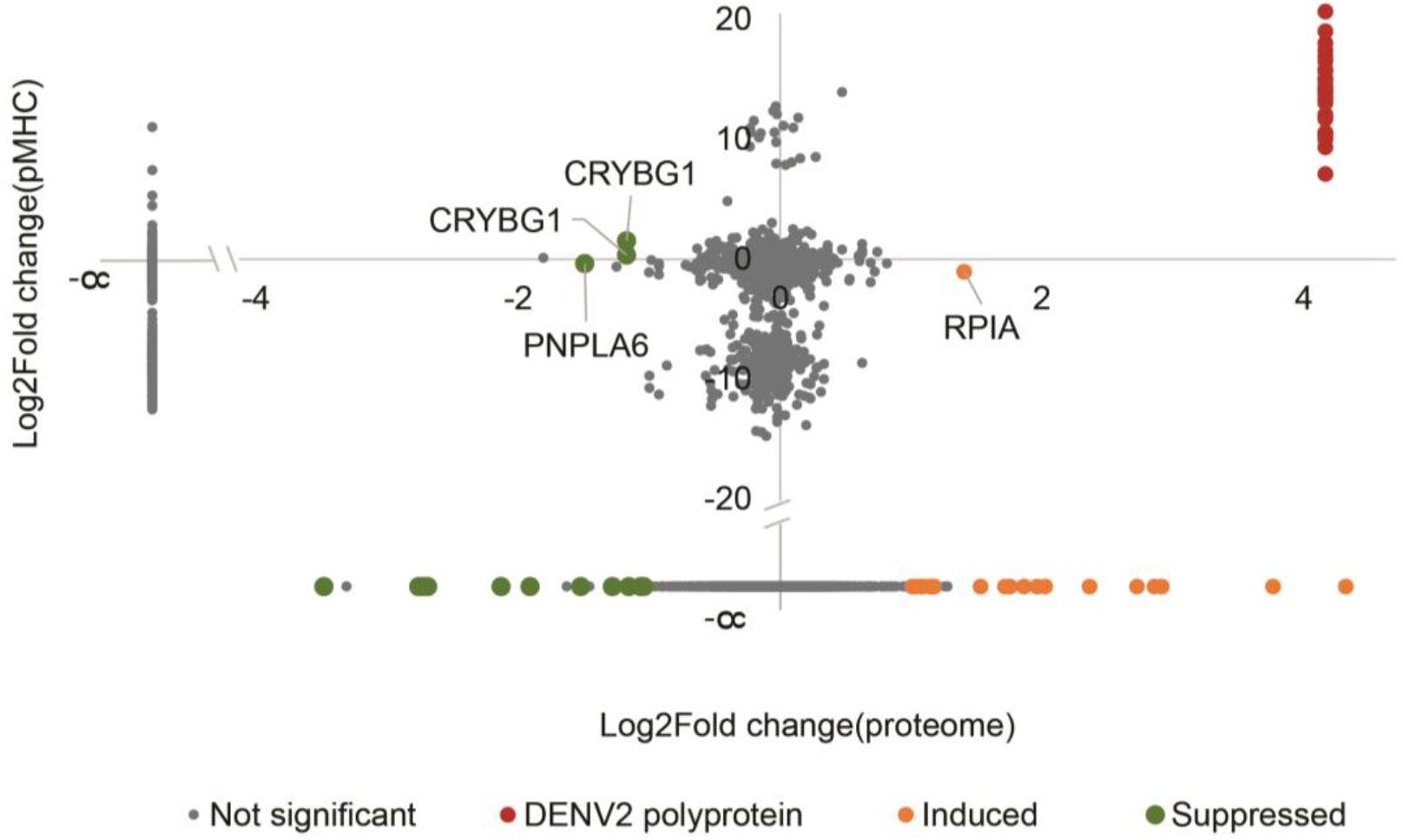
Proteome changes poorly predict pMHC repertoire shifts. Log2 fold change of proteins upon infection calculated from TMT reporter ion abundances in the proteome (x-axis) versus changes in their corresponding pMHCs. pMHCs from source proteins significantly (p < 0.01) suppressed (Log2 FC <1) or induced (Log2 FC > 1) upon DENV infection are highlighted in green and red respectively. DENV-derived pMHCs are highlighted in red.

**Supplementary Figure 5.**
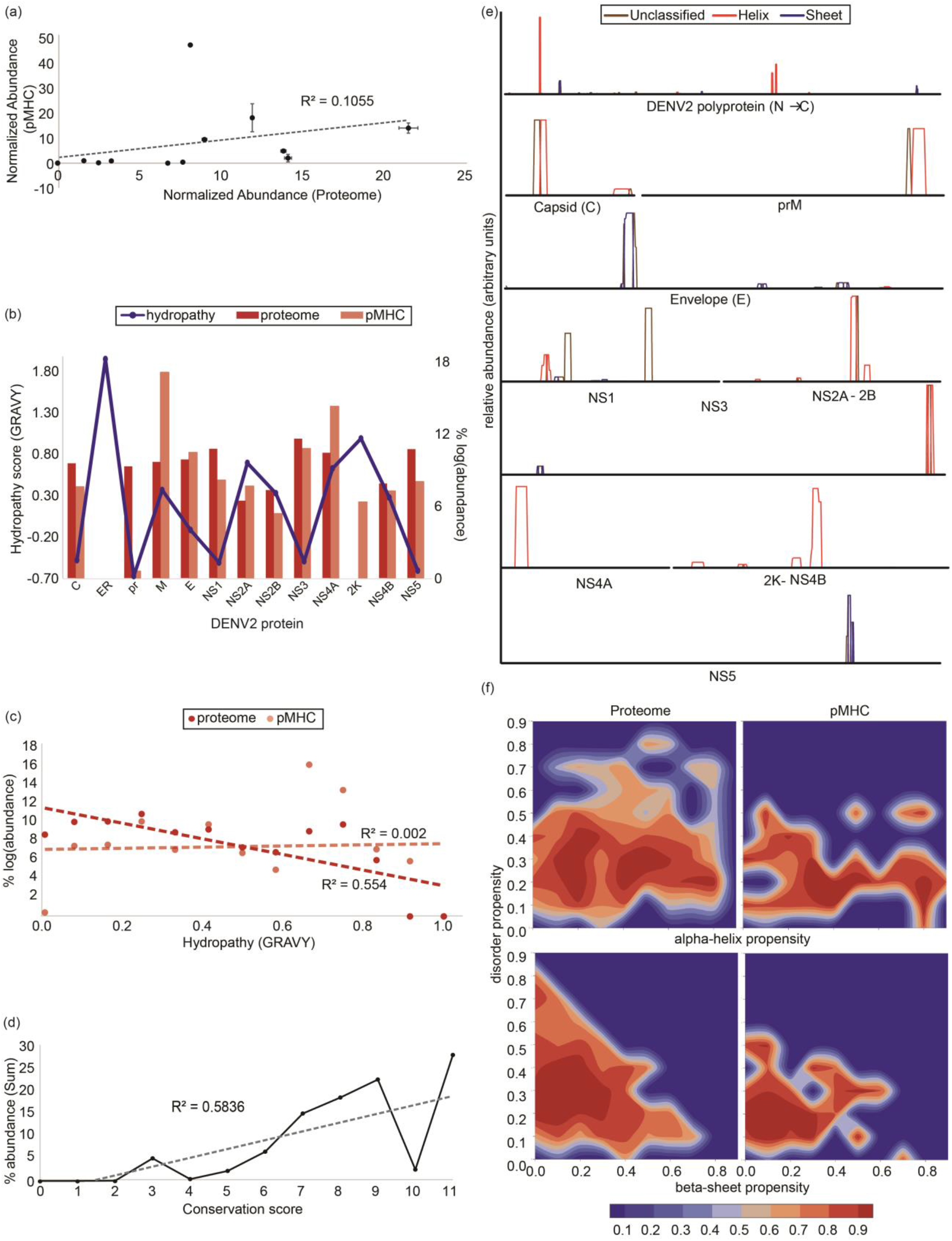
Structural features of the DENV polyprotein impact the viral pMHC repertoire. **a)** pMHC normalized abundance (y-axis) plotted against the source protein abundance (x-axis) in the tryptic proteome of infected cells normalized to account for number of tryptic cleavage sites. This plot suggests that viral source protein abundances poorly correlate with corresponding pMHC levels. **b)** Proteome (dark red) and pMHC (light red) abundance (y-axis, right) of each DENV polyprotein component (x-axis) and their predicted hydropathies represented as the GRAVY score (see methods) (blue) (y-axis, left) shows that protein hydropathies do not sufficiently explain protein or pMHC levels measured by LC-MS. **c)** Scatter plots of pMHC (orange) and proteome (red) hydropathies (x-axis) and their normalized log-transformed abundance quantify the modest or lack of correlation between protein hydropathies and their proteome or pMHC abundances respectively. **d)** Correlation between residue conservation using the AAcon score on the Jalview platform and pMHC abundance shows that residue conservation correlates modestly with presentation propensity. **e)** Distribution of ?-helix and ?-sheet derived pMHCs (71 in this study) across the DENV polyprotein was from the reported secondary structure features in the UniProt entry of a polyprotein homolog (see methods) and suggests a bias towards presentation of alpha helix structures versus beta sheets. Y-axis represents relative abundance of peptides across each polyprotein component **f)** Contour plot contrasting the density (abundance) of alpha helices (top) and beta sheets (bottom) in the DENV polyprotein (right) and the pMHCs (left).

**Supplementary Figure 6.**
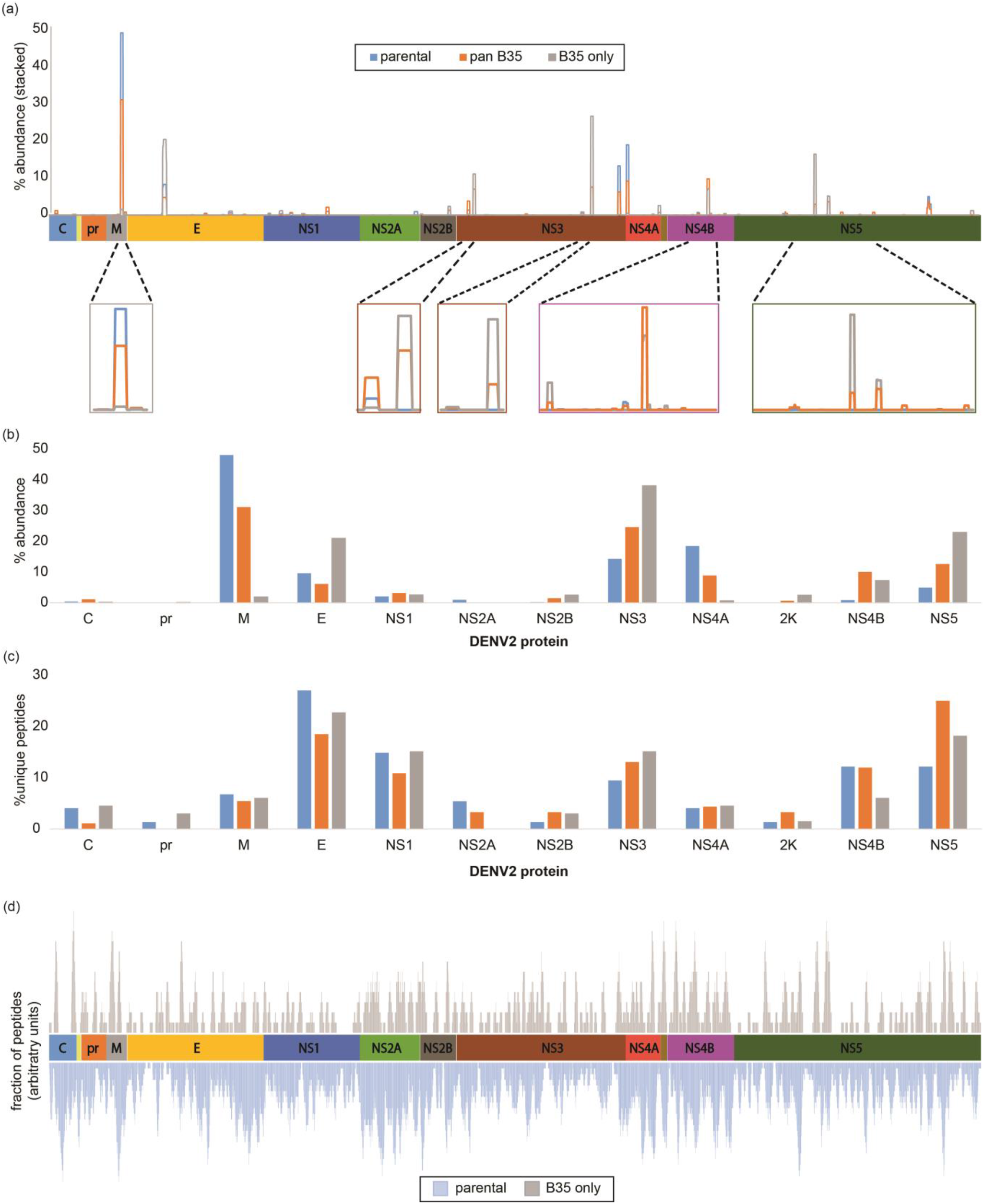
**a)** DENV pMHCs of parental (blue), B*35+endogenous (orange) Raji cells and B*35-only (grey) pMHCs mapped across the DENV polyprotein. Y-axis represents the stacked relative abundance of peptides spanning each residue. **b)** Percentage distribution of summed unique pMHCs from parental (blue), B*35+ endogenous Raji cells (orange) compared to the B*35-only pMHCs (grey) for each viral protein. **c)** Percentage distribution of summed pMHC abundance from parental (blue), B*35+ endogenous Raji cells (orange) compared to the B*35-only pMHCs (grey) for each viral protein. **d)** All 9-11 mer peptides predicted by netMHC to bind (< 5000 nM) B*35 (grey) to Raji HLAs are plotted below (light green) to reveal predicted binding hotspots. Y-axes represent relative number of peptides deemed binding spanning any given residue.

**Supplementary Figure 7.**
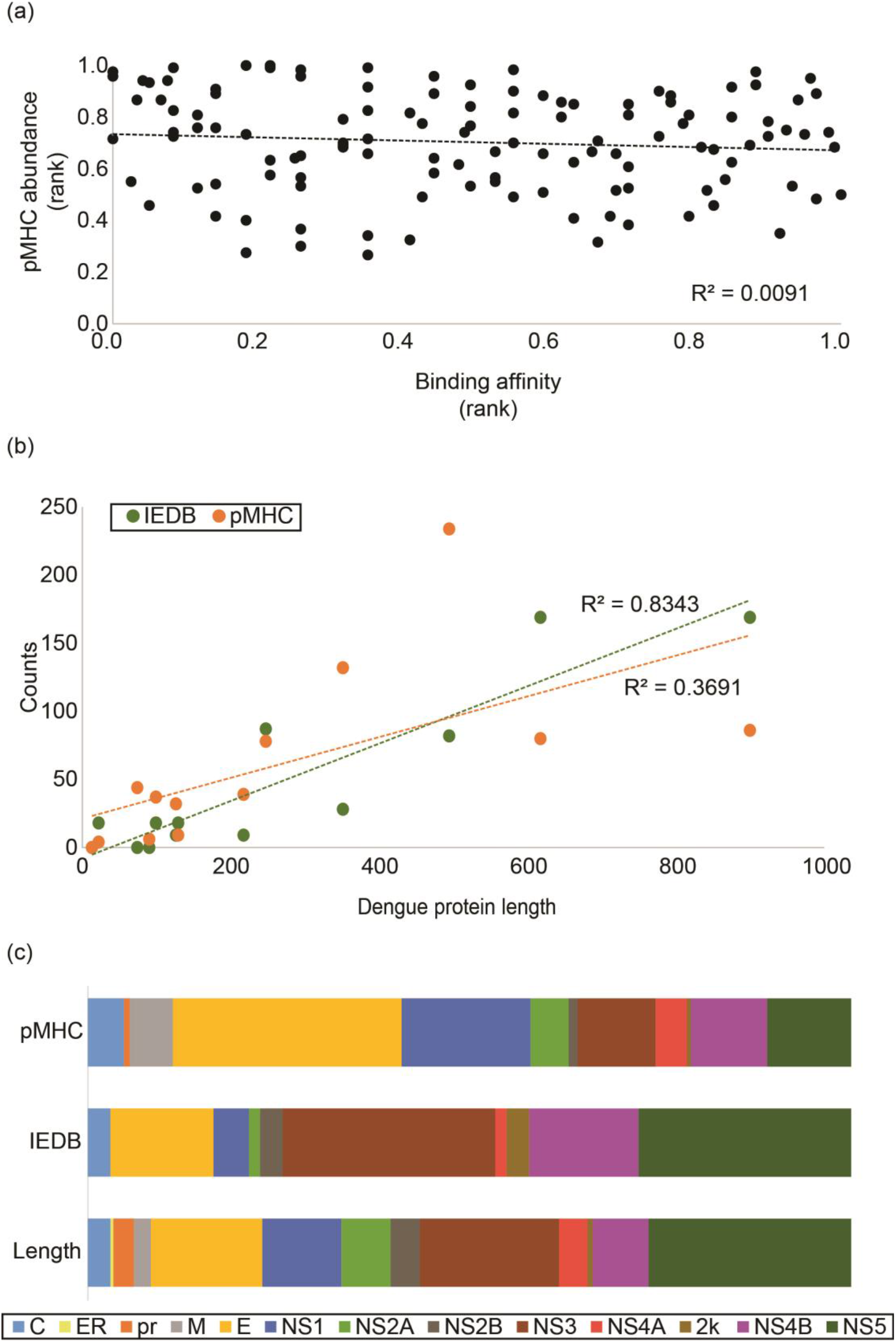
HLA-binding affinity is a poor predictor of presentation propensity. **a)** Correlation between DENV pMHCs’ predicted binding affinity (x-axis) and pMHC abundances (y-axis). Binding affinities represent percentile ranks of the dissociation constants for each pMHC (nM, lowest value for Raji endogenous HLA). Relative abundances were normalized to total DENV pMHC and percentile ranks were calculated. The correlation coefficient is indicated **b)** Correlation between DENV protein lengths (x-axis) and number of unique corresponding pMHCs (y-axis) from parental Raji cells (orange) and epitopes in IEDB (green). Correlation coefficients are indicated for each dataset. **c)** Stacked columns (to 100%) representing relative protein length (top), pMHC abundances (middle) and unique IEDB epitopes (bottom) summed protein-wise for DENV proteins.

### Supplementary Table 1

(a) List of all proteins in the TMT experiment

(b) Significantly modulated pathways inferred from proteome

### Supplementary Table 2

(a) List of Induced and Suppressed self-epitopes in DENV infected Raji cells (FC, p-value, IPA pathway information);

(b) All pMHC from control and DENV infected cells.

(c) Significantly modulated pathways inferred from pMHC data

### Supplementary Table 3

List of all DENV epitopes isolated – columns to indicate (1) all the experiments they were isolated in, (2) predicted (or experimentally established) restriction, (3) Empirical binding data - A*03 and B*35 MHC and (4) Average conservation of residues across DENV peptides

### Supplementary Table 4

Frequency of response against for every peptide across tested samples including positive (CMV), negative (HIV) controls as tested in ELISpot and tetramer staining assays.

### Experimental Procedures

#### Virus Stock

DENV-2 infectious clone 16681 was a gift from K. Kirkegaard. DENV-2 from infectious clone 16681 was adapted to HAP1 cells through serial passaging. Viral whole-genome sequence analysis revealed three coding mutations compared to the original clone 16681: Q399H in the envelope protein (E), L180F in NS2A and S238F in NS4B.

#### Plasmid constructs and genetic transductions

The cDNA sequence of full-length HLA-B*35:01:01:01:01 was synthesized (IDT) with 25 bp of overlapping vector sequence on either end and cloned into pLenti-CMV-Puro-DEST (w118-1, a gift from Eric Campeau) at the EcoRV sites using Gibson assembly (NEB). A two-step PCR was used to insert an N-terminal FLAG tag downstream of the signal peptide. Lentivirus produced in HEK293FT-cells was used to transduce Raji cells overnight. Transduced cells were selected by treatment with puromycin (1 μg/ml, InvivoGen) for 7 days.

#### Cell culture and viral infection

A B-lymphocyte cell line (Raji cells) overexpressing the viral entry receptor DC-SIGN (gift from Dr. Eva Harris, UC Berkeley) with and without HLA-B*35:01 overexpression were cultured in T175 flasks in RPMI medium supplemented with 5% Fetal Bovine Serum and 1x Penicillin/Streptomycin and L-glutamine, in two replicate experiments. The cells expanded to achieve 5e^8^ cells. One-half (2.5e^8^ cells) of these were infected with DENV-2 infectious clone 16681 at MOI of 5 and co-harvested at 27 hpi with the control cells. The harvested cells were washed twice with 1xPBS, flash frozen in liquid nitrogen and stored at −80°C until use.

#### Flow cytometry analysis

Harvested Raji cells were washed in 1xPBS, fixed with 4% paraformaldehyde for 10 mins at room temperature. The cells were washed again and stored in 1xPBS at 4°C until further analysis. For the detection of DENV NS1, cells were permeabilized in methanol for 30 mins at −20°C and then washed in FACS buffer (PBS, 2% FBS, 1 mM EDTA). The primary antibodies used were an anti-MHC class I (clone W6/32, Genentech), mouse IgG2α isotype control (Biolegend), anti-NS1 (Abcam), mouse IgG1 isotype control (Biolegend) and anti-FLAG (Sigma). Goat anti-mouse IgG-AlexaFluor647 (LifeTech) was used as a secondary antibody. The MHC staining was performed for 20 mins at 4°C at 1/100 dilution with 1/100 goat serum. The secondary antibody was used at 1/1000 for 20 mins at 4°C. The DENV anti-NS1 antibody was used 1/50 with 1/100 goat serum for 105 mins on ice followed by 1/1000 secondary and 1/100 goat serum for 1 hr on ice. All FLAG staining was performed at an anti-FLAG dilution of 1/100 and rat serum for 20 mins at room temperature and 1/1000 secondary with 1/100 rat serum for 20 mins at room temperature.

#### Multiplexed whole cellular proteome analysis

Multiplexed quantitative analysis of the whole cell proteome was carried out using isobaric tandem mass tag (TMT) labeling of two replicate samples of control and DENV infected parental Raji cells. A portion of the harvested cells (1e^7^ cells) from each sample were lysed for 15 mins in a bath sonicator on ice using a buffer (1 mL/1e^7^ cells) containing 8 M Urea, 100 mM NaCl and 25 mM Tris (pH 8.2), supplemented with Protease and Phosphatase inhibitor cocktail tablets (Roche). The cell debris was pelleted by centrifugation at 2500xg at 4°C and soluble protein was collected from the supernatant and quantified using the BCA protein assay kit (Thermo Scientific).

Protein disulfide bonds were reduced with dithiothreitol (5mM final concentration) at 56°C for 30 mins and alkylated with iodoacetamide treatment (15mM final concentration) for 1 hr in the dark. Protein digestion was carried out overnight at 37°C using MS-grade trypsin (Promega, 1:200 w/w) following dilution of Urea concentration to 1 M. The digest was quenched using 0.5% v/v trifluoroacetic acid and the peptides were desalted using C18-solid phase extraction and vacuum dried in a centrivap coupled to a cold-trap (Labconco).

Multiplexed TMT quantification was carried out as previously described^59^. Briefly, 100 μg of dried peptides from each sample were resuspended in 50 mM Na-HEPES buffer (pH 8.5) containing 30% anhydrous Acetonitrile. TMT reagents 126, 127N, 127C, 128N were added to tryptic peptides from control/DENV infected parental Raji samples and incubated for 1hr at room temperature. The reactions were quenched with 0.3% v/v hydroxylamine and then acidified with formic acid to achieve a pH of 2. A small, equal volume (2 μL) of each sample was combined and assessed by mass spectrometry to establish reporter ion intensity ratios. Adjusted amounts (where applicable) of the samples were mixed to achieve a 1:1:1:1 ratio, purified by C18 solid phase extraction and dried down. Dried peptides were fractionated by high pH reverse phase (HPRP) chromatography on an offline Agilent 1200 HPLC system using a C18 Extend column (Agilent). The 96 fractions collected were pooled as previously described^60^ to result in 12 concatenated samples, which were dried down and purified by C18 solid phase extraction. Purified peptides were stored at −80°C prior to LC-MS analysis. Four other samples were included in the initial TMT labeling and LC-MS analysis for a total of eight labeling conditions per result file, although data from these were not used or presented in this study.

#### Preparation of cell lysates for immunoprecipitation

Frozen cell pellets from uninfected (control) and DENV infected Raji cells were thawed on ice and lysed with buffer (1 mL/1.25×10^8^ cells) containing 1% CHAPS, 20 mM Tris and 150 mM NaCl (pH 8), supplemented with protease inhibitor cocktail (Roche), HALT (1x final concentration) and PMSF. Lysis was performed by incubating on ice for 20 mins with gentle vortexing every 5 mins. The cell lysates were transferred into 1.5 mL tubes (Eppendorf) and centrifuged at 16,000xg for 20 mins at 4°C to remove cellular debris and the supernatant was pre-cleared for 1 hr at 4°C using Protein-A Sepharose beads (GE Healthcare).

#### MHC-I peptide complex (pMHC) immuno-precipitation

Immunoprecipitation of the pre-cleared supernatant was performed using a pan-MHC-I antibody (W6/32, Genentech) coupled to Protein-A sepharose beads, on a rotating platform for 12 hrs at 4°C. The captured pMHC-I were eluted from the Protein-A Sepharose beads using 10% acetic acid and filtered using a 10 kDa cut-off filter to separate the peptides from the MHC-I molecules. Eluted peptides were vacuum dried in a centrivap coupled to a cold-trap (Labconco), re-suspended in 5% formic acid and desalted by binding to the C18 resin for solid phase extraction as previously described^63^.

#### B*35 FLAG immunoprecipitation

Anti-FLAG antibody-magnetic bead complexes (Sigma) prepared per manufacturer’s instructions, were added to clarified cell lysates and incubated overnight on a Hula mixer (Thermo) at 4°C. Unbound protein from the lysate was washed off the beads using 1xTBS at room temperature on a Hula mixer. The captured B*35 pMHC-I were eluted from the anti-FLAG beads using 10% acetic acid and filtered using a 10 kDa cut-off filter to separate the peptides from the MHC-I molecules. Eluted peptides were vacuum dried, resuspended in 5% formic acid and desalted as described above.

#### Liquid chromatography coupled with tandem mass spectrometry

De-salted MHC peptides from the W6/32 and FLAG pulldowns and from TMT experiments were re-suspended in sample buffer containing 0.1% Formic acid. Samples were separated by capillary reverse-phase chromatography on an 18 cm reversed-phase column (100 μm inner diameter, packed in-house with ReproSil-Pur C18-AQ 3.0 m resin (Dr. Maisch)) over a total run time of 160 min using a two-step linear gradient with 4–25% buffer B (0.2% (v/v) formic acid, 5% DMSO, and 94.8% (v/v) acetonitrile) for 120 min followed by 25–40% buffer B for 30 min using an Eksigent ekspert nanoLC-425 system (SCIEX, Framingham, Massachusetts, USA).

Three injections were made per MHC peptide sample to utilize multiple fragmentation modes (HCD (higher-energy collisional dissociation) or CID (collision-induced dissociation)). The third injection was performed with CID including singly charged species. MS data were acquired in data-dependent mode with the full MS scans collected in the Orbitrap mass analyzer with a resolution of 60,000 and m/z scan range 340 – 1,600.

The top ten most intense ions were then selected for sequencing and fragmented in the Orbitrap mass analyzer at a resolution of 15,000 (full width at half maximum). Data-dependent scans were acquired from precursors with masses ranging from 700 to 1,800 Da. Precursor ions were fragmented with a normalized collision energy of 35% and an activation time of 5 ms for CID and 30 ms for HCD. Repeat count was set to 2 and fragmented m/z values were dynamically excluded from further selection for a period of 30 s. The minimal signal threshold was set to 500 counts.

For TMT-labeled peptides, full MS scans were acquired in the Orbitrap mass analyzer with resolution 60,000 at 340 – 1,600 m/z. Unassigned charge states were rejected and the top 20 most intense ions with charge states >2 were sequentially isolated for MS/MS analysis using CID fragmentation. A minimal signal of 500 was required, the normalized collision energy was set at 35%, and the fragmented peptide masses were collected in the ion-trap. Dynamic exclusion was enabled with a repeat count of 1 and the repeat duration set to 30 s. MS2 fragment ions were further subjected to HCD fragmentation with multinotch MS3^62^ to yield the reporter ions from the TMT reagent which were then analyzed in the orbitrap of the mass spectrometer.

The mass spectrometry proteomics data have been deposited to the ProteomeXchange Consortium via the PRIDE^64^ partner repository with the dataset identifier PXD010280.

#### Generation of custom proteome sequence databases

A custom database was built comprising sequences from the human proteome UniProtKB including Swiss-Prot and TrEMBL databases (version May 2015), translated DENV-2 genome sequence (clone 16681) and common contaminant protein sequences included (for example, Staphylococcus protein A). The DENV polyprotein genome sequence was translated into its corresponding protein sequence using the ExPASy translate tool (https://web.expasy.org/translate/) from the Swiss Institute of Bioinformatics. Reversed ‘decoy’ protein sequences were appended to the database for ‘target-decoy’ error estimation^65^.

#### Analysis of cellular proteomes Tandem Mass Tag (TMT) data

TMT data were analyzed with ProteomeDiscover version 2.1.0.81 (Thermo Fisher Scientific) and SEQUEST-HT with the following settings: the parent mass error tolerance was set to 20 ppm. and the fragment mass error tolerance to 0.6 Da. Strict trypsin specificity was required, allowing for up to two missed cleavages. Carbamidomethylation of cysteine was set as fixed modification and oxidation of methionines as a variable modification. The minimum required peptide length was set to seven amino acids. All spectra were queried against the human database containing DENV and contaminant sequences as described above. A false discovery rate of 1% was required at both the peptide level and the protein level, calculated as the q-value by the Percolator algorithm^66^. For each protein group, summed peptide reporter ion intensities were used to estimate total protein abundance.

#### Computational identification of HLA peptides from mass spectra

All tandem mass spectra were queried against the custom database described above using both SEQUEST (version28.12)^67^ and PEAKS DB search engines (PEAKS Studio 7.5, Bioinformatics Solutions)^68^. Spectra were also interpreted by *de novo* sequencing (PEAKS Studio 7.5, Bioinformatics Solutions) to improve high-confidence peptide identification. The ms-convert program (version 3.0.45) was used to generate peak lists from RAW data files, and spectra were interpreted with SEQUEST. RAW data files were directly imported into PEAKS Studio 7.5 and subject to default data refinement (deisotoping, charge deconvolution, peak centroiding) prior to searching with PEAKS DB and PEAKS *de novo* algorithms. For all searches, the parent mass error tolerance was set to 20 ppm. and the fragment mass error tolerance to 0.02 Da. For SEQUEST and PEAKS DB, enzyme specificity was set to none and oxidation of methionines and deamidation (N, Q), cysteinylation, and phosphorylation (S, T, Y) were considered as variable modifications.

High-confidence peptide identifications were selected at a 1% false discovery rate with a modified version of the Percolator algorithm^66^, optimized for proteogenomic immunopeptide analysis as previously described^36^.

#### Comparisons between MHC presentation and protein expression levels

pMHC relative abundances were inferred from peak areas of corresponding peptides and normalized to the total area signal for each run. Peptides reproducibly measured in both biological replicates were used to determine which self-pMHCs were significantly altered upon infection. Log2 transformed peptide areas were compared using paired two-tailed t-tests (Qlucore Omics Explorer) to determine pMHCs significantly (p<0.01) upregulated or downregulated during DENV infection. Missing data points were deemed below detection limit and imputed as the minimum value in that dataset. pMHCs were ranked in their decreasing order of relative abundance in both control and infected systems and rank changes (ΔRank) were compared^37^ to measure which peptides accounted for the differences between the two states.

Protein abundances in the cellular proteome were inferred from reporter ion intensities in the TMT experiments. Abundances were weighted by the predicted number of tryptic cleavage sites to infer normalized protein abundance. Proteins that significantly changed during infection were determined using Qlucore Omics Explorer as described for the ligandome above.

Protein and ligandome log2-transformed fold-changes were compared to determine if protein expression levels directly impacted MHC-presentation. Ingenuity Pathway Analysis was used to determine cellular pathways significantly (-log10 p-value > 1.3, righttailed Fisher’s Exact test) perturbed during DENV infection using the log-transformed fold change and p-values associated with the proteins in the proteome dataset. Pathway-level changes reflected in the ligandome data were inferred in a similar manner using the log2-transformed fold change and p-value associated with the ligandome data.

#### Structural characterization of proteins and peptides

pMHCs derived from the DENV polyprotein were mapped on to the most homologous template available in the UniProt database - DENV-2 strain Thailand/16681/1984 (accession P29990) to infer sequence features. Secondary structure and disorder propensities of host- and DENV-derived peptides in the ligandome and proteome datasets were calculated using VSL2^69,70^ and PSIPRED^71^. In-house wrapper scripts were used to run these programs and to assign the peptide disorders. Peptides were assigned to 10 bins according to the helix or disorder propensity scores assigned by PSIPRED. Visualization of these data as contour plots of helix vs. disorder propensities was implemented in Plotly (https://plot.ly) using a custom R-script.

The tertiary structure of DENV polyprotein was predicted using homology modeling on the SwissModel (swissmodel.expasy.org) platform hosted on the Expasy server (Swiss Institute of Bioinformatics). Structures were visualized and annotated on Jmol (www.jmol.org).

Peptide hydropathies were calculated as the length-normalized grand average of hydropathy (gravy-calculator.de) on the Sequence Manipulation Suite^72^.

Multiple Sequence Alignment of all complete DENV serotypes (1-4) sequences on UniProt was performed using BLASTp^73^. Jalview^74^ was used to visualize aligned sequences and calculate the conservation score for each residue across the DENV polyprotein.

#### Epitope-HLA *in vitro* binding assays

Classical competition assays to quantitatively measure peptide binding to HLA A*03:01 and B*35:01 class I MHC molecules were based on the inhibition of binding of a high-affinity radiolabeled peptide to purified MHC molecules. MHC purification and binding assays performed as detailed elsewhere^75^. Briefly, 0.1 – 1 nM of radiolabeled peptide was co-incubated at room temperature with 1 μM to 1 nM of purified MHC in the presence of a cocktail of protease inhibitors and 1 μM β2-microglobulin. Following a two-day incubation, MHC-bound radioactivity was determined by capturing MHC/peptide complexes on W6/32 (anti-class I) antibody coated Lumitrac 600 plates (Greiner Bio-one, Frickenhausen, Germany), and measuring bound counts per minute (CPM) using the TopCount (Packard Instrument Co., Meriden, CT) micro-scintillation counter. In the case of competitive assays, the concentration of peptide yielding 50% inhibition of the binding of the radiolabeled peptide was calculated. Under the conditions utilized, where [label] < [MHC] and IC50 ≥ [MHC], the measured IC50 values are reasonable approximations of the true Kd values^76,77^. Each competitor peptide was tested at six different concentrations covering a 100,000-fold dose range, and in three or more independent experiments. As a positive control, the unlabeled version of the radiolabeled probe was also tested in each experiment.

#### Ex vivo IFN-γ ELISPOT assays

PBMCs were prepared from laboratory-confirmed DENV-seropositive donors from the Nicaraguan National Blood Bank and the Colombo National Blood Bank (SriLanka) as previously described^78^. PBMCs (2×10^5^ cells/well) were incubated in triplicates with 0.1 mL of complete RPMI 1640 (Omega Scientific) supplemented with 5% human serum (Cellgro) in the presence of HLA-matched peptide pools (2 μg/mL), as previously described^47^. Briefly, following 20 hr incubation at 37°C, the cells were incubated with biotinylated IFNγ mAb (mAb 7-B6-1; Mabtech) for 2 hrs and developed as previously described^47^. Phytohemagglutinin (PHA) and A*03 and B*35 restricted CMV epitopes were used as positive control and HIV (A*03 and B*35) epitopes were used as negative controls.

#### HLA-A*03 and B*35 tetramer staining and preparation

HLA-A*03:01 and B*35:01 tetramers containing an ultraviolet-cleavable peptide^79^ were synthesized by the NIH Tetramer Facility. Seventeen DENV peptides were synthesized (ELIM Biopharmaceuticals). Potential HLA-A*03 or B*35-binding (predicted IC50 < 500 nM by netMHC3.4) binding peptides with A*03- and B*35-monomers were exchanged, and multimerized with streptavidin-PE, APC, PECy7 or BV605 as previously described^80^. Peptides that bound both HLA-A*03 and HLA-B*35 with poor affinities were exchanged and multimerized to generate both A*03 and B*35 tetramers. PBMCs from three HLA-A*03 and four HLA-B*35 individuals (mutually exclusive) seropositive for DENV-2 were donated by Alessandro Sette at the La Jolla Institute for Allergy and Immunology. To determine background staining, we used T-cells from leukocyte reduction system chambers from a healthy untyped donor from the Stanford Blood Center. Tetramers generated from IEDB B*35 restricted peptides were pooled to allow multiplexing and address low donor cell numbers. Tetramer staining was performed as previously described^80^. HIV peptides RLRPGGKKK and NSSKVSQNY were used as A*03 and B*35 negative controls respectively, and CMV peptides TTVYPPSSTAK and IPSINVHHY were used as A*03 and B*35 positive controls.

